# Mechanosensory stimulation triggers sustained local motor activity in *Drosophila melanogaster*

**DOI:** 10.1101/2022.07.19.500315

**Authors:** Alexandra M. Medeiros, Anna F. Hobbiss, Gonçalo Borges, Marta Moita, César S. Mendes

## Abstract

Most vertebrates and invertebrates such as Drosophila melanogaster are able to move in complex environments due to their ability to integrate sensory information along with motor commands. Mechanosensory structures exist along each leg to assist in motor coordination by transmitting external cues or proprioceptive information to motor centers in the central nervous system. Nevertheless, how different mechanosensory structures engage these locomotor centers and their underlying circuits remains poorly understood.

Here, we tested the role of mechanosensory structures in movement initiation by optogenetically stimulating specific classes of leg sensory structures. We found that stimulation of leg Mechanosensory Bristles (MsB) and femoral Chordotonal Organ (ChO) is sufficient to initiate forward movement in immobile animals. While the stimulation of the ChO required brain centers to induce forward movement, unexpectedly, brief stimulation of leg MsB triggered sustained cyclic motor activity dependent only on circuits within the Ventral Nerve Cord (VNC). The duration of the MsB-induced movement was dependent on the number of excited cells and specific to leg afferents, since stimulation of MsB in other segments lead to different motor outcomes. MsB-mediated movement lacked inter and intra-leg coordination, but preserved antagonistic muscle activity within joints. Our data shows that sensory stimulation can act in combination with descending commands in order to elicit a faster response to mechanical stimulation. In addition, it sheds light on the ability of specific sensory circuits to modulate motor control, including initiation of movement, presenting a new system to better understand how different levels of coordination are controlled by VNC and central brain locomotor circuits.

**Significance Statement:** Sensory feedback is critical to allow smooth and stable locomotion. Proprioceptors interact directly with pre-motor centers optimizing and sustaining coordinated movement. However, initiation of moment is considered to be triggered by higher-order centers in the brain. Here we took advantage of the genetic toolkit provided by the fruit fly Drosophila melanogaster to optogenetically activate different classes of leg sensory cells in immobile animals. We found that leg mechanosensory bristles can specifically trigger sustained leg activity independently of higher-order centers as headless flies could sustain prolonged leg movement upon mechanosensory stimulation. Moreover, while this sensory-evoked movement lacks intra- and inter-leg coordination, it still preserved basic antagonistic muscle activity. These findings suggest a parallel mechanism to trigger fast movement upon sensory stimulation. In addition, it provides a new model for movement initiation and a point–of-entry to define pre-motor circuits.

## Introduction

Moving organisms display remarkably complex and yet highly efficient motor circuits capable to integrate multisensorial information and execute the appropriate motor response (reviewed by (1, 2) (3)). In *Drosophila*, descending interneurons (DNs) act like “command-neurons” bridging decision centers in the central brain with executive circuits in the Ventral Nerve Chord (VNC), the insect equivalent of the mammalian spinal cord, eliciting a plethora of motor behaviors including locomotion (4–8). Equally remarkable, is the ability of these circuits to quickly adapt to external environmental conditions, optimizing speed, stability and energy consumption (1, 2, 9). Key for this flexibility is mechanosensory feedback, including proprioception and exteroception, enabled by specialized neurons that relay internal and external physical features such as muscle extension, tissue compression or gravitational orientation to motor centers. Lack of sensory feedback renders animals highly uncoordinated, largely determined by the number and class of inactivated cells (10, 11). Sensory structures vary in shape and properties depending on the type of mechanical feature being transduced (12). For example, in insects, the Chordotonal organ (ChO) relays the mechanical features of joint angles, a structure analogous to mammalian muscle spindles (13), whereas Campaniform Sensilla (CS), dome-like structures in the cuticle of the fly leg, sense deformation caused by load (13, 14).

Another crucial mechanosensory structure are the Mechanosensory Bristles (MsB). These structures are involved in detecting external cues, including dust particles, parasites and wind intensity, triggering varied motor outcomes such as grooming, defensive kicking or locomotion arrest (15–18). In the context of collective behavior, mechanosensory bristles play a role in conspecific avoidance by initiating walking responses triggered by another fly’s touch (19). An avoidance response was also seen after the optogenetic stimulation of mechanosensory bristles in specific legs during walking (20). Given that self-generated movement also results in bristle deflection, these structures may also play a role in proprioception(13).

While dendrites of these mechanosensory neurons are located in the leg segment containing the sensory cell, axons project into the central nervous system, mostly into the ventral-most layer of the leg neuropil within the VNC, where sensory information is processed (13, 16, 21–23). Sensory structures interact with local circuits in the VNC in order to modulate and adjust locomotor patterns. However, to generate fully functional and coordinated locomotion, communication between motor circuits and higher order centers is also required (24). Evidence shows that locally these structures can directly affect ongoing muscle contraction in order to correct movement and posture. For instance, ChO receptors that sense the position of the femur-tibia joint react to an imposed extension of the tibia by inducing excitation of flexor tibiae motor neurons and at the same time activate local interneurons that inhibit the antagonistic extensor tibiae motor neurons (25). Another example of local modulation by sensory inputs comes from the stick insect, where it was shown that sensory feedback, notably from the ChO and campaniform sensilla, assists in phase transition from stance to swing phase, a process termed “active reaction”, which influences the activation threshold by flexor and extensor motor pools providing a positive reinforcement of movement (26, 27). Further studies suggest the existence of a local neuronal architecture within the VNC linking particular sensory structures to Central Pattern Generators (CPGs) and leg motor neurons to produce inter-segmental coordination (13, 28). These features allow the generation of a continuous, stable and coordinated walking pattern facilitating step transitions. These data also raise the possibility that proprioception can trigger movement initiation. However, whether fly circuits physically substantiate this possibility, and whether distinct sensory neuron types differ in their ability to activate movement, remains unknown. To investigate this question, we took advantage of the rich *Drosophila* genetic tool-kit and quantitative kinematic tools available for this model.

Here we show that stimulation of MsB or ChO is sufficient to initiate forward movement from an immobile fly. Interestingly, we found that while leg ChO stimulation depends on the presence of a central brain for flies to engage in forward movement, brief leg MsB stimulation could trigger an immediate and sustained motor activity independently of central brain activity. This response is also specific to leg MsB since stimulation in the wing or head lead to different motor outcomes. The MsB-evoked response can be triggered from different leg segments and its intensity depends on the number of stimulated bristles. Leg MsB-triggered forward movement lacks inter and intra-leg coordination although antagonistic muscle activity is nevertheless preserved.

This study sheds light on the ability of specific sensory circuits to engage motor circuits in the ventral nerve cord independently of descending commands. Moreover, it suggests how and what aspects the central brain can control during forward coordinated locomotion.

## Results

### Activation of leg mechanosensory bristles evokes central brain independent movement

To understand the role of leg sensory afferents in movement control and initiation we identified fly lines that targeted the different types of mechanosensory neurons present in the leg. For that, we performed an expression screen for GAL4 lines from the Janelia Flylight Collection that labelled specific sensory structures in the fly leg (29, 30). In order to restrict GAL4 expression to the legs, we used a combinatorial approach with flipase under the control of the Dac^RE^ enhancer fragment (11). Among the relevant lines and based on the expression pattern we selected 2 lines, one labelling Mechanosensory Bristles (hereafter termed MsB1) and one labelling the femoral Chordotonal Organ (termed ChO1) (Fig. 1A, B, and S1). Although normally classified as exteroceptors, MsB can additionally be described as proprioceptors since these afferents can also be activated during self-movement, which is supported by the observation that their inactivation causes perturbations in walking patterns (13) (Medeiros and Mendes, unpublished). Moreover, based on the type of sensilla associated with dendritic structures and their axonal profiles we ruled out that no chemosensory structures were labeled (16, 31, 32). We chose ChO line due to the large body of work focusing on this structure (12, 13, 15, 33–38).

**Figure 1.**
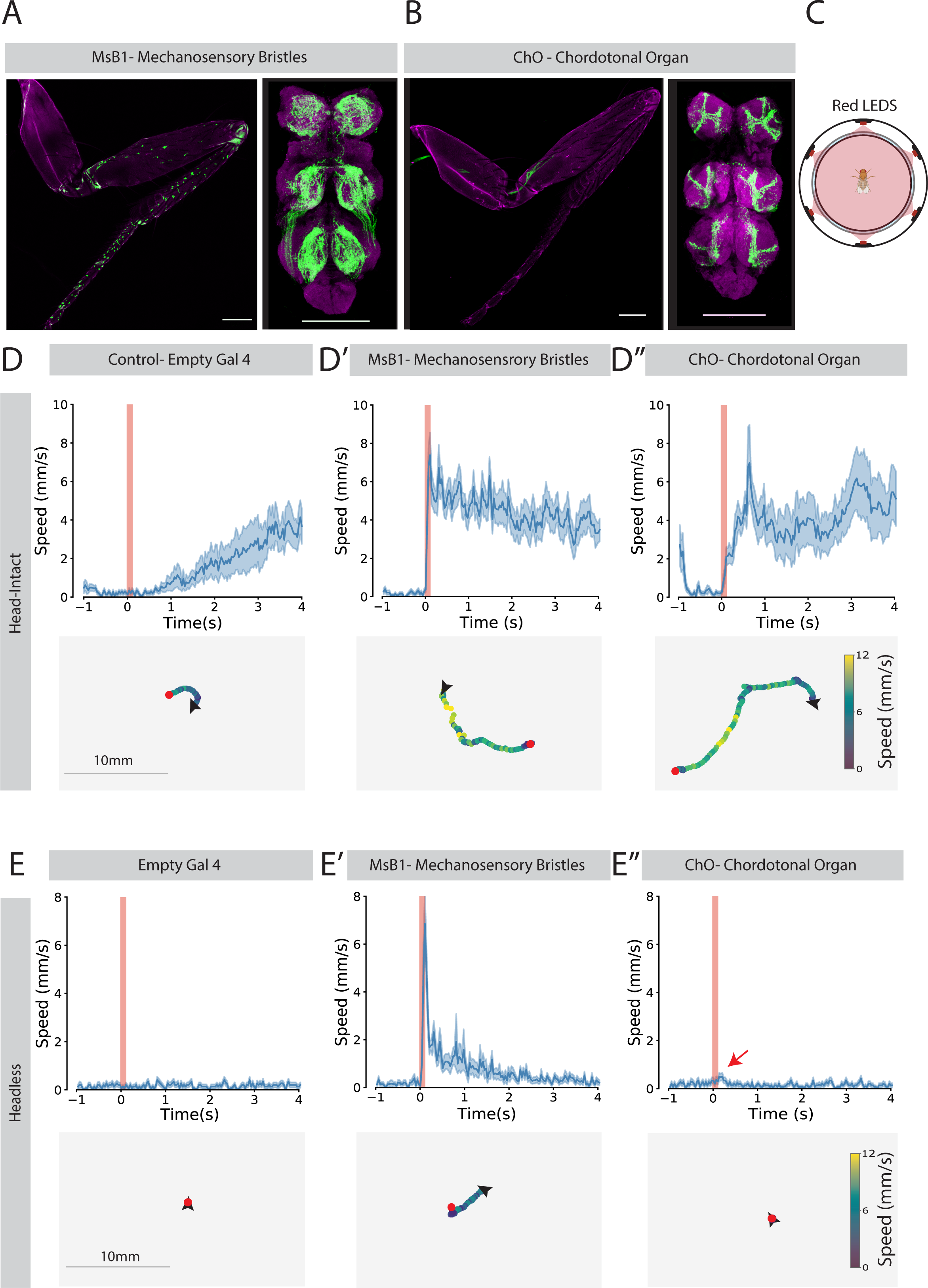
Optogenetic stimulation of leg mechanosensory bristles triggers movement in stationary flies. (A) Leg and VNC expression pattern of leg MsB. mVenus expression under combinatorial control of R65D12 and dac^RE^-flp. Genotype: R65D12-GAL44, dac^RE^-flp, UAS-FRT-stop-FRT-Chrimson-mVenus. Cell bodies are located in the legs (left panel) and send axonal projections to the VNC (right panel). Bar, 100 μm. (B) Leg and VNC expression pattern in femoral ChO neurons. mVenus expression under combinatorial control of R79E02 and ato-flp. Genotype: R79E02-Gal4, ato-flp, UAS-FRT-stop-FRT-Chrimson-mVenus. Cell bodies are located in the legs (left panel) and send axonal projections to the VNC (right panel). Bar, 100 μm. (C) Schematic of the behavioral arena. Arena is surrounded by 6 deep red LEDs. See Methods for details. (D, E) Instantaneous speed and trajectories over time after optogenetic stimulation. Upper panels show instantaneous speed over time with red bar representing 100 ms red light stimulus. Dark line and light blue shadows indicate the average and SEM for all replicates, respectively. Lower panels show representative trajectories for 4 seconds after optogenetic stimulation. Trajectory plots are color coded according to the speed of the fly with black arrows and red dots indicating the initial and final positions, respectively. (D) Head intact and (E) headless animals. Red arrow indicates a brief speed increase due to reflexive leg twitch. N= 20 animals for all conditions, except for group MsB head-intact N=19.

We tested the expression patterns of the selected lines in sensory neurons by visualization of mVenus tagged to the light-gated channel CsChrimson used in subsequent experiments (39). We observed that as described previously, MsB1 neurons project to the ventral-most layer of the three leg Neuropils, the ventral association center (VAC) (23) (Fig. S1A). Conversely, the medial ventral association layer (mVAC) and the intermediate Neuropil (IntNP) received projections from ChO1 line neurons (23) (Fig. 1 and S1B). In the central brain we identified ChO1 axon terminals in the Wedge (WED) and AMMC consistent with the previously identified ChO primary neuron projections to the central brain (Fig. S1B). The MsB1 line showed some labelling in the Antennal lobes and GNG, possibly axon terminals from head appendages (16, 40–42). After close inspection of the head appendages we found cell labelling in the maxillary palps, in the proboscis and in the 3^rd^ antennal segment (Fig. S1A and data not shown).

Next, we measured the behavioral responses of immobile flies to optogenetic stimulation of these 2 classes of sensory neurons. For this we drove the expression of the light-gated CsChrimson channel in the respective structures and analyzed the flies’ behavior after a single 100 ms stimulation in an open field arena (see Methods). While control Empty-GAL4 flies remained immobile immediately after stimulation, eventually resuming motor activity, the stimulation of leg mechanosensory Bristles or the leg femoral ChO led to movement (Fig. 1D and Video S1). Motor responses differed between these two lines, with MsB1 stimulation inducing a very short latency response, reaching a maximum speed of 7.38 ± 1.17 mm/s, 60 ms after stimulus onset, while ChO1 stimulation led to a more progressive response reaching maximum speed of 6.98 ± 1.99 mm/s after 600 ms (Fig. 1D’ and D’’, upper panels). Moreover, after careful inspection of the videos we noted that MsB1 stimulation triggered a fast startle response characterized by an apparent uncoordinated locomotor pattern followed by sustained coordinated walking (Fig. 1D’, Video S1). On the other hand, ChO1 stimulation led to an apparent coordinated walking pattern (Fig. 1D’’, Video S1).

The different responses to optogenetic stimulation between ChO and MsB stimulation suggested that these neurons promote movement by different neuronal circuits. We then asked if movement initiation was dependent on descending information, by decapitating the animals before optogenetic stimulation (Fig. 1E). Decapitated Drosophila are capable of maintaining an upright posture and display innate behaviors such as grooming (43, 44).

Control animals (empty-GAL4) did not exhibit any optogenetic response, maintaining an upright posture with occasional grooming behavior (Fig. 1E and data not shown). Strikingly, MsB1 stimulation triggered a startle response and sustained motor activity for approximately 2.9 seconds (Fig. 1E’ and Video S1). Moreover, under these conditions, animals seem to phenocopy the uncoordinated behavior observed in head-intact animals, reaching a peak speed of 6.85 ± 1.27 mm/s, 60 ms after stimulus onset, suggesting that the initial response to mechanosensory stimulation is mediated by circuits located within the VNC, independent of descending circuits. In contrast, ChO1 stimulation in beheaded flies did not induce movement initiation (Fig. 1E’’), indicating that these sensory neurons activate central brain circuits that in turn are responsible for walking initiation. Optogenetic stimulation of ChO1 in decapitated flies did however lead to a reflexive leg twitch, which induced a small increase in the perceived instantaneous speed (red arrow in Fig. 1E’’and Video S**1**). This reflex is most likely the consequence of direct synaptic connections between the ChO and leg motor neurons, as described in locusts (13, 25, 45, 46).

These results indicate that stimulating mechanosensory structures in the leg is sufficient to induce walking initiation, which in the case of MsB only depends on local circuits within the VNC. However, sustained walking after MsB stimulation requires descending activity from the head.

### Cyclic movement triggered by MsB is leg specific

Having shown that stimulation of MsB is capable of triggering movement independently from descending commands we asked whether this motor response can also be triggered by MsB present in other body structures. Full body stimulation of mechanosensory bristles (using dust particles or genetically) has been shown to trigger grooming behavior (18, 43), while in our study where we limit activation to just leg MsB, we found mostly uncoordinated walking to be the main response. Considering the previous findings indicating that compartmentalized sensory connections mediate different behavioral responses (44, 47), we asked if the motor response could be determined by the body structure from which the stimulus originates. To target expression of CsChrimson to MsB in other body segments, we used the same GAL4 driver but instead combined with a head capsule or wing-specific flipase line using the eyeless (ey) and vestigial (vg) fragments, respectably (Fig. 2A, see methods). We confirmed the effectiveness of this approach based on the presence of positive cells in the respective cuticle and axonal projections (Fig. S1C, D and E). We then cataloged the repertoire of motor behaviors in response to optogenetic stimulation using automatic and manual classifiers including uncoordinated and coordinated movement, leg cycling and grooming (see Methods).

**Figure 2.**
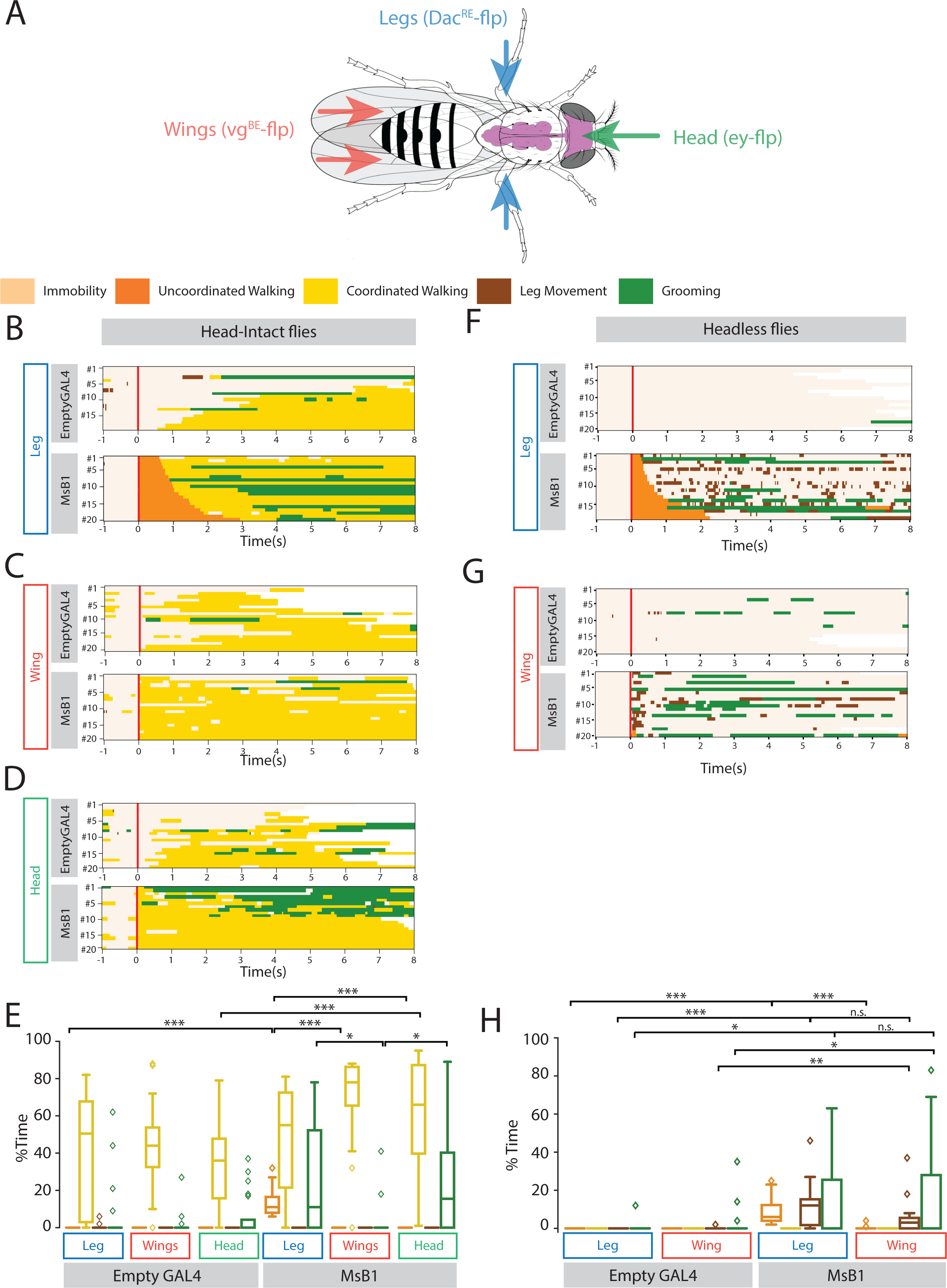
Cyclic movement is specific to leg MS bristle stimulation. (A) Schematic showing the combinatorial genetic tools used to restrict the expression of the MsB GAL4 driver R65D12 to the legs (DAC^RE^-flp), wings (Vg^BE^-flp) and head (ey-flp) regions. Each combination was crossed with UAS>>CsChrimson. (B-D, F-G) Raster plot showing the type of movement performed by the flies before and after the stimulation of leg MsB. X axis represents the time in seconds where 0 is the moment of the red light pulse stimulation (red Bar). Each line corresponds to one individual. Movement categories: Immobility (beige); Uncoordinated walking (orange); Coordinated walking (yellow); Leg movement (brown); and Grooming (green). Upper Panels shows Empty GAL4 behavior while lower panel shows MSB1 GAL4 (R65D12) behavior (B-D) Head-intact animals. (B) Leg domain (with DAC^RE^-flp). (C) Wing domain (with Vg^BE^-flp). (D) Head domain (with ey-flp). (E, H) Percentage of time spent doing one of the behaviors described for (B-D,F-G). Boxplots represent the median as the middle line, with the lower and upper edges of the boxes representing the 25% and 75% quartiles, respectively, whiskers represent the range of the full data set, excluding outliers represented by diamonds. Statistical analysis with Kruskal–Wallis-ANOVA followed by Dunn’s *post hoc* analysis, *p<0.05 **p<0.01, ***p<0.001. (E) Quantification for head-intact animals as shown in (B-D). (F-G) Headless flies. (F) Leg domain (with DAC^RE^-flp). (G) Wing domain (with Vg^BE^-flp). (H) Quantification for head-intact animals as shown in (F-G). N= 20 for all conditions.

In head-intact animals, leg MsB activation triggered an apparent uncoordinated displacement, immediately followed by coordinated walking and grooming in most animals (12 out of 20) (Fig. 2B). Notably, wing and head stimulation did not induce uncoordinated movement, with animals instead initiating coordinated walking (Fig. 2C-E), showing that the uncoordinated activation of leg motor centers is specific to leg afferents. Moreover, stimulation of wing MsB led only a small fraction of the animals to groom (2 out of 20) compared to leg or head stimulation (11 and 10 out of 20, respectively) (Fig. 2B-E), further highlighting the specificity of sensory stimulation for motor selection. This suggests that behavior selection (in this case grooming) is dependent on the place where the stimulus is received.

We then asked if the aforementioned behaviors are dependent on the presence of higher-order brain commands. Leg MsB stimulation in headless animals caused an apparent uncoordinated body displacement (Fig. 2F and H). Additionally, close inspection of the movies revealed an extra behavior, which is interestingly absent in head intact animals: sustained random leg movements (Figs. 2F, brown sections and Video S2). These movements do not display any synchrony between legs, thus rendering the fly immobile.

Our data also show that the walking behavior triggered by the stimulation of wing MsB is dependent on the presence of higher-order brain commands. Stimulation of wing MsB afferents in headless flies did not induce walking or uncoordinated movement (Fig. 2G, H), but instead grooming, (a behavior which is absent in head intact conditions (Fig. 2C), suggesting a descending inhibitory command that blocks the grooming behavior). Video analysis of headless flies showed that the observed leg movement bouts resembled the preparatory pose for grooming (data not shown), or in some cases a defensive kick behavior described previously to remove fictitious parasites (17) (Video S2).

Overall, our data show that the motor response triggered by sensory stimulation depends not only on the class of proprioceptors (MsB), but also on the source of the stimulus. These data also highlight the role of the central brain controlling VNC pre-motor centers, while executing particular leg motor commands such as leg twitching or grooming.

### Mechanosensory-triggered movement is dependent on the number of cells

Mechanosensory bristles are present on the surface of the entire leg of the fly, with neurons present in the most distal part of the leg projecting their axons to the central part of the leg neuropil while neurons occupying proximal positions project to more peripheral areas (16). Leg MsB display a concentric organization in the leg neuropil instead of a concentrated topographical distribution within the VNC (16). Since we showed that an apparent uncoordinated movement is evoked specifically by leg MsB, we conjectured the possibility that a discrete subpopulation of leg cells could be sufficient to activate the downstream motor program that triggers cyclic movement, or alternatively, that the response results from the contribution of all leg MsB being activated. To test these possibilities, we used 6 additional GAL4 lines to drive the expression of CsChrimson in different bristles subpopulations in the fly leg, which we called MsB2 to 7 (with our original line termed MsB1), ordered by the number of cells labeled, with MsB1 the most and MsB7 the least (Table S1). For each line we quantified number of MsB labeled and looked at their localization in specific leg segments (Fig. 3A, B and Table S1). We also measured the response of the different lines to optogenetic stimulation by registering the duration of motor activity and categorizing the different motor outputs in an open field arena (Fig. 3C and D, respectively). As previously described (Fig. 1) the MsB1 line, which expressed GAL4 in 169 bristles in all leg segments, had a strong reaction to the stimulus, showing motor activity for up to 1.5 minutes (Fig. 3C). Interestingly, stimulation of a lower number of cells in MsB2-6, which label between 57-26 neurons, still triggered a motor response but with a shorter time span (Fig. 3C), whilst MsB7, which drives expression in just 4 cells in the tarsal segments, triggered minimal motor activity, suggesting that the duration of movement is influenced by the number of activated cells. To further analyze the type of behaviors triggered by mechanosensory bristles stimulation, we used the same behavioral classifier previously used to identify immobility, uncoordinated walking, grooming, and sporadic leg movement (Fig. 3D, E and S3). We observed that most lines show uncoordinated walking behavior followed by sporadic leg movement without displacement. Some flies also displayed grooming behavior, consistent with previous observations (43) (Fig. 3D, E).

**Figure 3.**
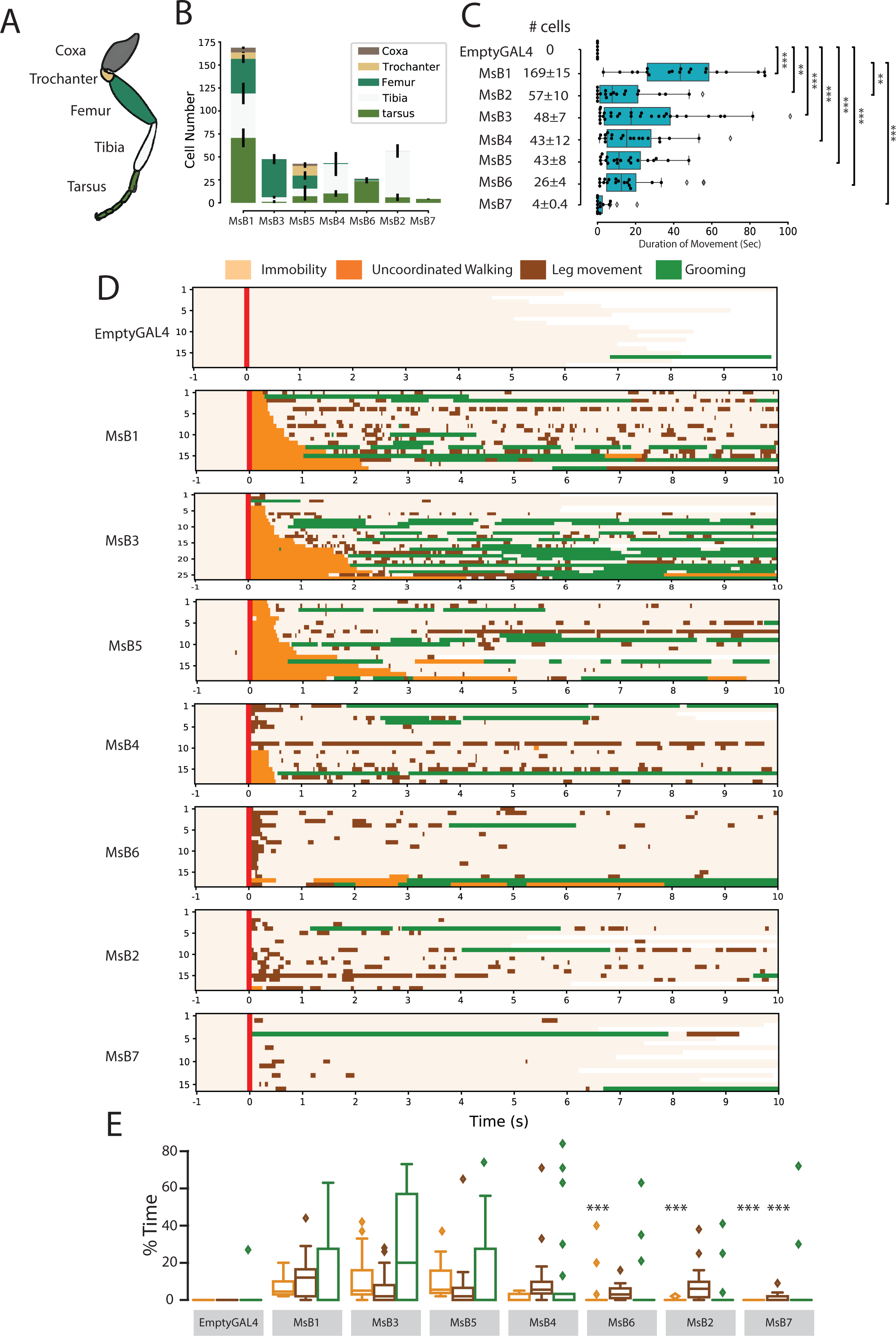
Movement pattern triggered by distinct leg MsB populations in headless animals. (A) Schematic depicting the leg segments of the of *Drosophila melanogaster*. (B) Cell labelling distribution amongst leg segments for each line. Error bar represent standard deviation. MSB1: R65D12[leg] N= 20, MSB2:R77E09[leg] N=18, MSB3:R39A11[leg] N=27, MSB4: R54A11[leg] N=20, MSB5: R64D07[leg] N=20, MSB6: R85A07[leg] N=20, MSB7: R45D07[leg] N=17. (C) Duration of movement after one pulse of red light stimulation (100ms) for each MSB line. Lines are organized according to the number of labeled cells in an ascending order. (D) Raster plots showing the type of movement performed by the flies in each group, 1 second before and up to 10 seconds after the stimulation. X axis represent the time in seconds being 0 the time of optogenetic stimulation (red Bar). Each line corresponds to one individual. The categorization of movements includes Immobility (beige), Uncoordinated Walking (orange), Leg Displacement (brown); and Grooming (green). Lines are ordered in descending order according to the length of uncoordinated walking response (E) Boxplots showing for each MSB line the percentage of time spent doing a specific behavior. Behaviors are color-coded as in panel D. Boxplots represent the median as the middle line, with the lower and upper edges of the boxes representing the 25% and 75% quartiles, respectively, whiskers represent the range of the full data set, excluding outliers represented by diamonds. Statistical analysis with Kruskal–Wallis-ANOVA followed by Dunn’s *post hoc* analysis, *p<0.05, **p<0.01, ***p<0.001.

We then tested the influence of the number of MsB activated in the duration of the Uncoordinated phase only (Fig.3D). We did not find any correlation between the total number of labelled cells and the duration of the uncoordinated phase (r=0.17). Interestingly, by looking at the number of cells labelled in different segments we observed that the number of labeled cells in femur showed a positive correlation with the uncoordinated movement (r=0.44), while cells present in other structures: tarsus, tibia, trochanter and coxa, had lower or no correlation at all (r=0.06, r=0.13, r=0.28, r=0.21, respectively). These results suggest that no particular population of MsB is uniquely responsible for inducing VNC-dependent leg movement, although MsB located in the femur show a higher influence in the motor response. Furthermore, the cyclic motor response is due to the cumulative stimulation of multiple MsB broadly distributed circuits in the VNC.

### MsB stimulation evokes movement encompassing a non-coordinated phase

We next investigated the kinematic features associated with VNC-dependent motor response. For this we used the FlyWalker system that allows the quantification of kinematic parameters with high spatial and temporal resolution generating a large set of outputs including step, spatial and gait parameters (11, 36, 48). The walking chamber was modified in order to accommodate a far-red LED array to perform optogenetic stimulation where each fly received one pulse of 100 ms red light stimulation and the resulting behavior was quantified.

Control animals display a typical gait pattern showing an alternating tripod configuration (Fig 4A) and consistent stance traces (Fig 4C), which mark the tarsal contacts relative to the body axis during stance phases, indicating high levels of leg coordination under these conditions (11, 45, 49–51). Forward movement induced by leg MsB stimulation can be classified into two distinct phases using visual inspection of the step patterns: an early *uncoordinated phase* followed by a late *coordinated phase* (Fig. 4B/D and B’/E, respectively; and Movie S3). The *uncoordinated phase* immediately follows optogenetic stimulation, displaying an apparent random step pattern without any clear coupling between the six legs, leading to an inefficient force transduction that results in a slow and non-cyclic movement (Fig. 4B). We further analyzed the step configurations, a proxy for inter-leg coordination, and found a large proportion of frames that do not match any of the normal gaits or configurations described by control flies, such as tripod or tetrapod (11, 49, 50, 52) (Fig. 4B, lower panel). We describe these configurations as ‘*non-compliant*’ since these do not conform with the rules of coordinated movement described by Durr et al. (53). Non-compliant configurations include, for example, two contralateral or two consecutive ipsilateral legs lifted simultaneously, which creates static instability. We observed that non-compliant configurations are used to a striking degree during the uncoordinated phase, ranging from 20% to 50% (average of 29.6±4.4%) of the frames compared to being nearly absent in control animals (Fig. 2F and H). This shift in gait pattern is matched by a reduction in the use of the canonical tripod and tetrapod gaits (Fig. 4H). These data suggest that stimulation of leg MsB induces activation of motor activity lacking inter-leg coordination. Moreover, stance traces display an extremely uncoordinated pattern with highly variable and inconsistent leg contacts, highlighting a lack of control and coordination during this phase of locomotion (Fig. 4B and D), even though leg retraction shows the typical anterior to posterior movement (Fig. 4D). Nevertheless, the high degree of uncoordinated movement suggests that in addition to a lack of inter-leg coordination, the proper coupling between leg segments (or intra-leg coordination) is lost during this initial phase of locomotion. We evaluated the whole kinematic profile and found that virtually all kinematic parameters were altered, with the notable exception of the average swing duration (Fig. 4H). Interestingly, we found that animals shifted from an uncoordinated phase to a distinct coordinated phase without requiring a transitional stop (Fig. 4B and B’). Nevertheless, during the transition most flies displayed a period during which all legs were in a stance phase, contacting the ground (19 out of 21, corresponding to 90.4%) (Fig. 4B, ∼1.4s). From a total of 21 flies tested, 10 (48%) shifted to coordinated walking without stopping (see for example, Fig. 4B to 4B’), 4 (19%) shifted to coordinated walking after a small pause of 200 to 400 ms and 6 (29 %) flies initiated grooming for a significant period of time before resuming walking. During the coordinated phase the fly adopts mainly tripod and tetrapod gaits while non-compliance configurations are absent (Fig. 4B’, lower panel). Once animals have returned to a coordinated phase, both gait patterns and stance traces display a pattern similar to control animals (Fig. 4H, 4B’ and E). Kinematic parameters in the coordinated phase are more concordant with the standard walking pattern seen in control animals although the recovery is not total, indicating that transient sensory stimulation can have sustained effects on locomotion (Fig. 4H, 4B’).

**Figure 4.**
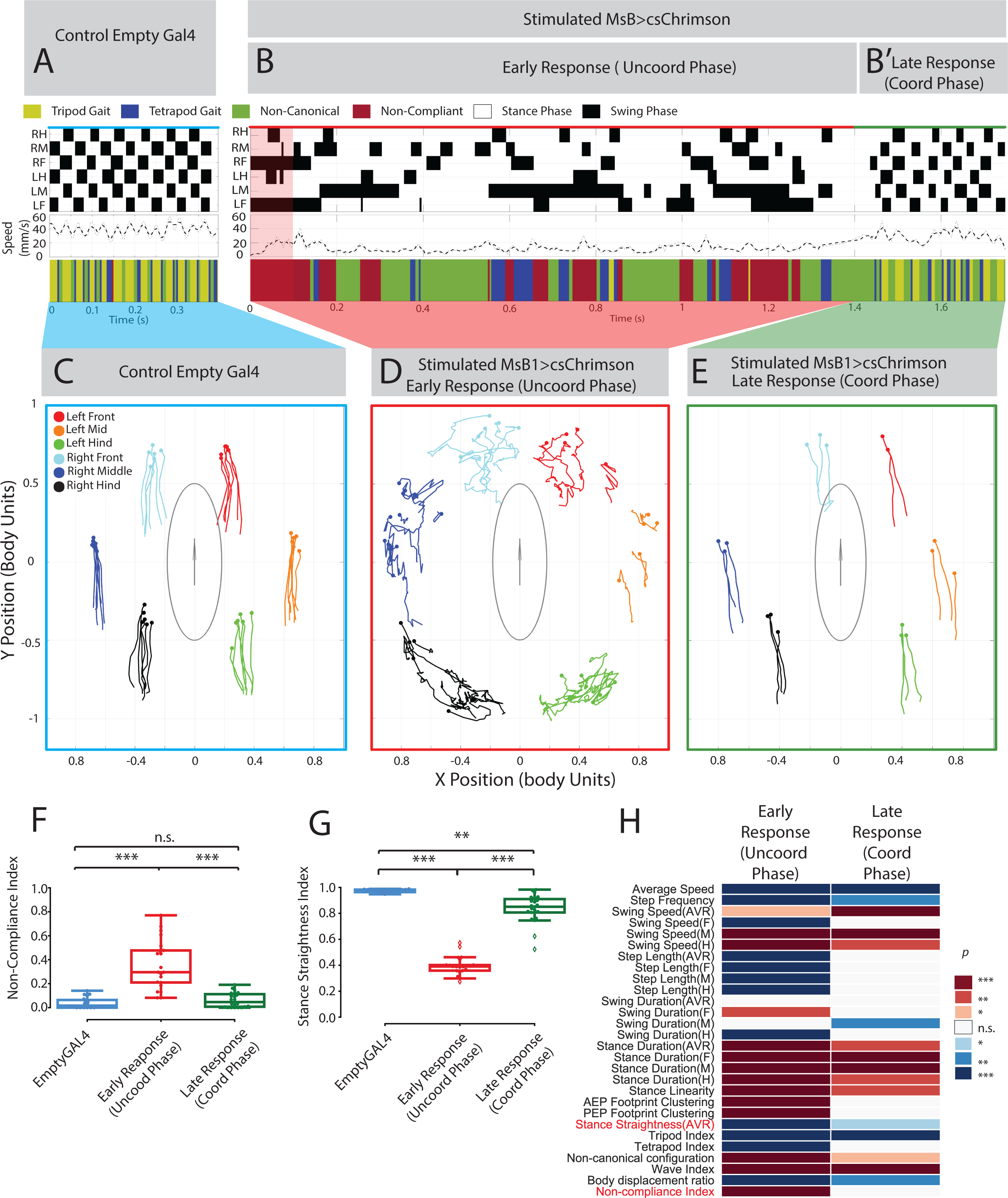
Kinematic analysis of walking patterns in MsB1 flies after mechanosensory bristles stimulation. (A-B) Representative gait features MsB-evoked movement. Upper panel shows the flies’ step patterns across time. For each leg, swing phases are represented in black and stance phases in white. Right hind (RH); right mid (RM); right front (RF); left hind (LH); left middle (LM); left front (LF). Middle panel shows the instantaneous speed across time for the same video. Thick and thin lines correspond to integration times of 25 or 12.5 ms, respectively. Lower panel represents the gaits adopted by the fly. For each frame the corresponding gait was color-coded as follows: Yellow (tripod), blue (tetrapod), green (non-canonical), red (non-compliant configuration). See Methods. 100ms optogenetic stimulation is represented by a red shadow. (A) Empty GAL4 control (Genotype: empty-GAL44, dac^RE^-flp, UAS-FRT-stop-FRT-Chrimson-mVenus). (B) Post-stimulus response of MSB1 flies (Genotype: R65D12-GAL4, dac^RE^-flp, UAS-FRT-stop-FRT-Chrimson-mVenus). 100ms optogenetic stimulation is represented by a red shadow. Sequence is divided in early response (Uncoordinated Phase) and late response (Coordinated Phase) (B and B’, respectively). (C-E) Representative *stance traces*. Traces are generated by the position of the stance phase footprints relative to the body center (set at 0.0,0.0). For each leg, stance onset corresponds to the *Anterior Extreme Position* (*AEP*) (colored circle) while stance offset is termed *Posterior Extreme Position* (*PEP*). Experimental conditions are color matched with panels (A-B). (C) Control Empty GAL4 control. (D) Post-stimulus of MSB1 flies, early response corresponding to the uncoordinated phase. (E) Post-stimulus of MSB1 flies, late response corresponding to the coordinated phase. (F-H) Kinematic quantification between the empty GAL4 control (N=20) and the early (N=21) and late response post stimulated (N=23). In (F-G) boxplots represent the median as the middle line, with the lower and upper edges of the boxes representing the 25% and 75% quartiles, respectively, whiskers represent the range of the full data set, excluding outliers represented by diamonds. Filled dots represent individual flies. Values are normalized for body size. Statistical analysis with Kruskal–Wallis-ANOVA followed by Dunn’s *post hoc* analysis, **p<0.01, ***p<0.001. (F) *Non-Compliance Index* (G) *Stance straightness index*. (H) Heat map of kinematic parameters comparing empty GAL4 control with early response/ Uncoordinated Phase (first Column) and late response/Coordinated Phase (second Column) of experimental group, with each line representing a motor parameter. Data were residual normalized and outliers were excluded. Statistical analysis with one-way ANOVA followed by Tukey’s post hoc test (for normal distributions) or Kruskal–Wallis-ANOVA followed by Dunn’s post hoc test (for non-normal distribution). p values are represented by a color code with red and blue shades indicate an increase or decrease relative to control, respectively. *P<0.05; **P<0.01; ***P<0.001. White indicates no statistically significant variation.

In summary, stimulation of leg MsB results in a reflexive forward cyclic movement, during which flies display an uncoordinated phase, lacking inter-leg coordination, followed by a coordinated phase presenting the stereotypical properties of walking animals.

### Kinematic features after leg MsB stimulation in headless flies

We next investigated the kinematic features of headless flies upon leg MsB stimulation (Fig. 5), which we previously found initiates locomotor behavior in the open arena setup (Fig. 1 E). As described for Figure 4, we optogenetically stimulated leg MsB1 in an enclosed arena and extracted kinematic features using the FlyWalker system (see Methods). Prior to stimulation, we decapitated the flies and glued the heads back to the thorax to allow proper body tracking by the FlyWalker software.

**Figure 5.**
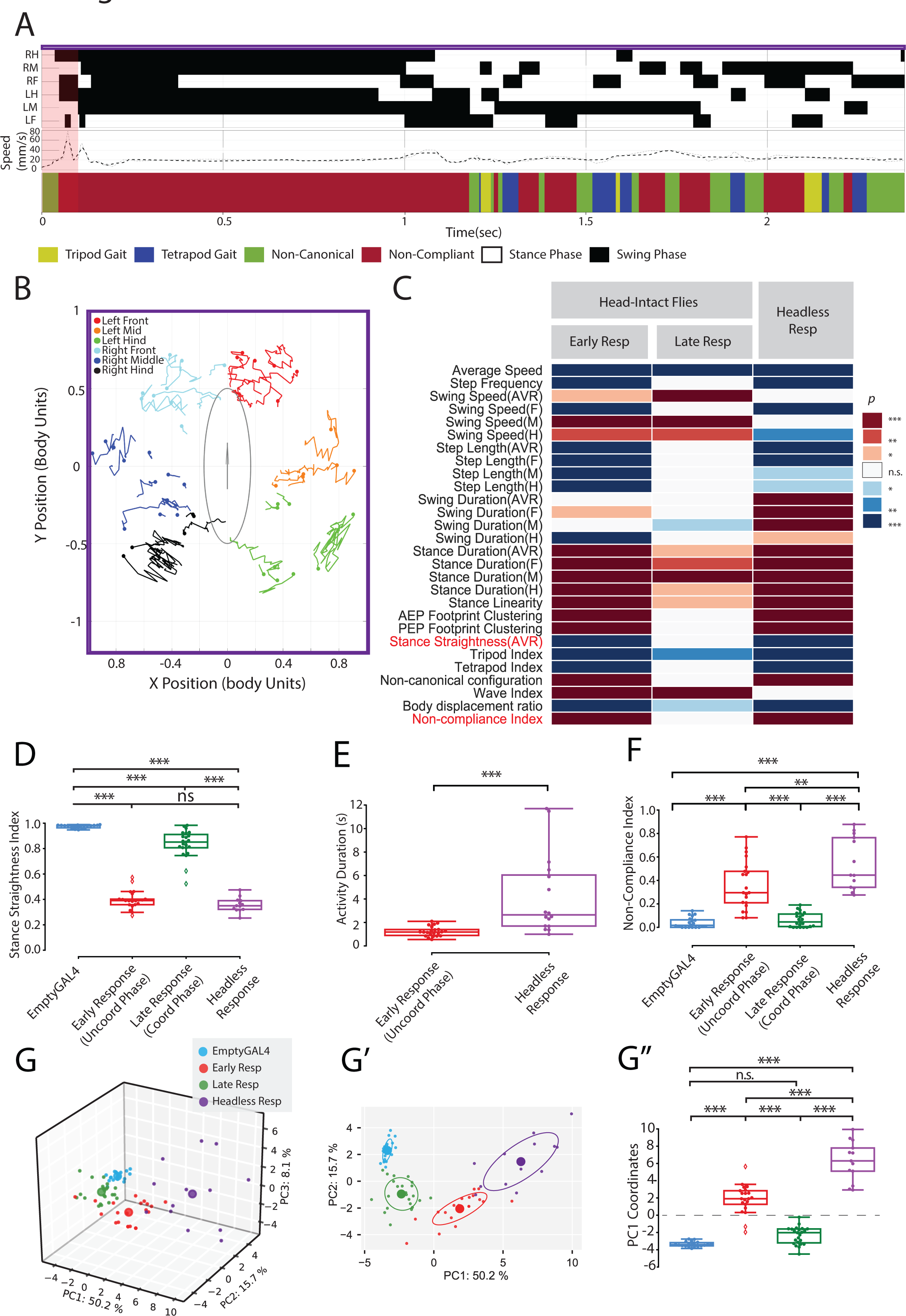
Mechanosensory Bristles evoked movement is independent of the central Brain. (A) Representative gait features after MsB stimulation of headless flies. Genotype: R65D12-GAL4, dac^RE^-flp, UAS-FRT-stop-FRT-Chrimson-mVenus. Upper panel shows the flies’ step patterns across time. For each leg swing phases are represented in black and stance phases in white. Right hind (RH); right mid (RM); right front (RF); left hind (LH); left middle (LM); left front (LF). Middle panel shows the instantaneous speed across time for the same video. Thick and thin lines correspond to integration times of 25 or 12.5 ms, respectively. Lower panel represents the gaits adopted by the fly. For each frame the corresponding gait was color-coded as follows: Yellow (tripod), blue (tetrapod), green (non-canonical), red (non-compliant configurations). 100ms optogenetic stimulation is represented by a red shadow. (B) Representative *stance traces* following optogenetic stimulation. Traces are generated by the position of the stance phase footprints relative to the body center (set at 0.0,0.0). For each leg, stance onset corresponds to the *Anterior Extreme Position* (*AEP*) (colored circle) while stance offset is termed *Posterior Extreme Position* (*PEP*). (C) Heat map of kinematic parameters comparing empty GAL4 control of head-intact animals (N=20) with the early/uncoordinated (N= 21) and late/coordinated responses (N=23) (first and second columns, respectively) of post-stimulated head-intact animals, and post-stimulated headless flies (N=13) (third column). Statistical analysis with one-way ANOVA followed by Tukey’s post hoc test (for normal distributions) or Kruskal– Wallis-ANOVA followed by Dunn’s post hoc test (for non-normal distribution). p values are represented by a color code with red and blue shades indicating an increase or decrease relative to control, respectively. *P<0.05; **P<0.01; ***P<0.001. White indicates no variation. (D) Stance straightness index in head-intact empty GAL4 controls, early and late response of post-stimulated head-intact animals, and post-stimulated headless flies. (E) Post stimulation activity between head-intact (early response) and headless flies. (F) Non-Compliance Index in head-intact empty GAL4 controls, early and late response of post-stimulated head-intact animals, and post-stimulated headless flies. (G-G’’) PCA of all kinematic parameters in head-intact empty GAL4 controls, early and late response of post-stimulated head-intact animals, and post-stimulated headless flies. (G) Tridimensional representation of three-component PCA analysis. Each individual small dot represents one video while larger dots represent the average point (n=20 for each condition). Contribution of each component is indicated in each axis. (G’) 2D representation of PC1-PC2 with ellipses delimiting 50% of variance of the data. (G’’) Comparison of each PC1 coordinates. In (D-F, G’’) boxplots represent the median as the middle line, with the lower and upper edges of the boxes representing the 25% and 75% quartiles, respectively, whiskers represent the range of the full data set, excluding outliers represented by diamonds. Filled dots represent individual flies. Values are normalized for body size. Statistical analysis with Kruskal–Wallis-ANOVA followed by Dunn’s *post hoc* analysis, *P < 0.05; **P < 0.01; ***P < 0.001. (D) PC1 coordinates. (E) PC1 coordinates. See Methods for details.

Stimulated flies display cyclic leg movements with a highly uncoordinated gait pattern and stance traces (Fig. 5A, B). Moreover, kinematic properties also display a starkly different profile compared to control animals (Fig. 5C). Noticeably, there is a similarity between the walking pattern of the uncoordinated phase of the head intact flies and the headless flies after stimulation (Fig. 5C). In fact, the stance straightness index is indistinguishable between these two conditions (Fig. 5D, compare Fig. 4D with 5B). However, we observed that the time headless flies spend moving after the stimulus is longer compared to the uncoordinated phase in head intact flies (Fig. 5E), suggesting that descending circuits can interrupt the uncoordinated cyclic activity triggered by leg MsB and mediate a transition to a coordinated walking pattern.

We then asked if descending control could influence the response to leg MsB stimulation by comparing the early response in head intact animals to the response in headless flies. First, we noticed a larger fraction of frames spent in a non-compliant configuration (Fig. 5F), which could be explained by a sustained swing phase during the initial stages of the motor response (Fig. 5A, below ∼1 sec, compare lower section, Fig. 4B with 5A). Second, we subjected all the kinematic parameters to a Principal Component Analysis (PCA) selecting the top 3 axes that explained most variance, thus yielding a three-dimensional space representation (Fig. 5G). PC1, which explained the highest variance of the data (50.2%), allowed a statistical discrimination between headless animals and the uncoordinated phase in head-intact animals, with the cloud of points of headless animals lying further away from control animals in the behavioral space (Fig. 5G’ and G’’), suggesting that higher order circuits can influence the cyclic motor activity triggered by leg MsB.

These data indicate that leg MsB can trigger sustained cyclic motor activity using solely circuits within the VNC, albeit with severely impaired coordination between legs and leg segments. This cyclic activity is nevertheless modulated and ultimately restrained by central brain circuits and descending information.

### Inter-joint coordination is partially disrupted after MsB stimulation

To further explore the disruption of coordinated locomotion following leg MsB stimulation, we examined intra-leg, or alternatively called inter-joint coordination, i.e. the correct modulation of joint angles within a single limb in relation to one another, required to produce efficient walking (54). To extract measures of joint angles we tethered flies expressing CsChrimson in leg MsB to a wire while they walked freely on a polystyrene ball (55–58) (Fig 6A). As above, one 100ms light stimulation pulse was delivered while the flies were immobile. Fly behavior was imaged using a lateral camera at 120 fps and videos were tracked using DeepLabCut (59), which allowed us to extract 3 joint angles: 1) Coxa-Femur joint, 2) Femur-Tibia joint, 3) Tibia-Tarsus joint (Fig. 6A, B and Video S4). We only considered angles from the T1 leg, since this leg is predominantly at a perpendicular angle to the imaging path and thus the angles could be accurately calculated in a 2D video. As expected from our earlier results, MsB stimulation induced movement in both head-intact and headless flies (Fig. 6B, Video S4). To determine whether inter-joint coordination was perturbed by the stimulation protocol we compared 4 conditions: walking flies (*pre-stimulation*) (Fig. 6B, top panel); head-intact flies for the initial 1s after stimulation onset (termed “*early response*”) (Fig. 6B, middle panel, red section); head-intact flies 3-5s after stimulation onset (termed “*late response*”) (Fig. 6B, middle panel, green section); and headless flies for the first 1s after stimulation onset (termed “*headless response*”) (Fig. 6B, lower panel, purple section). We chose these time periods to capture the ‘uncoordinated’ and ‘coordinated’ phases of post-stimulation movement previously described (Figs. 4 and 5).

**Figure 6.**
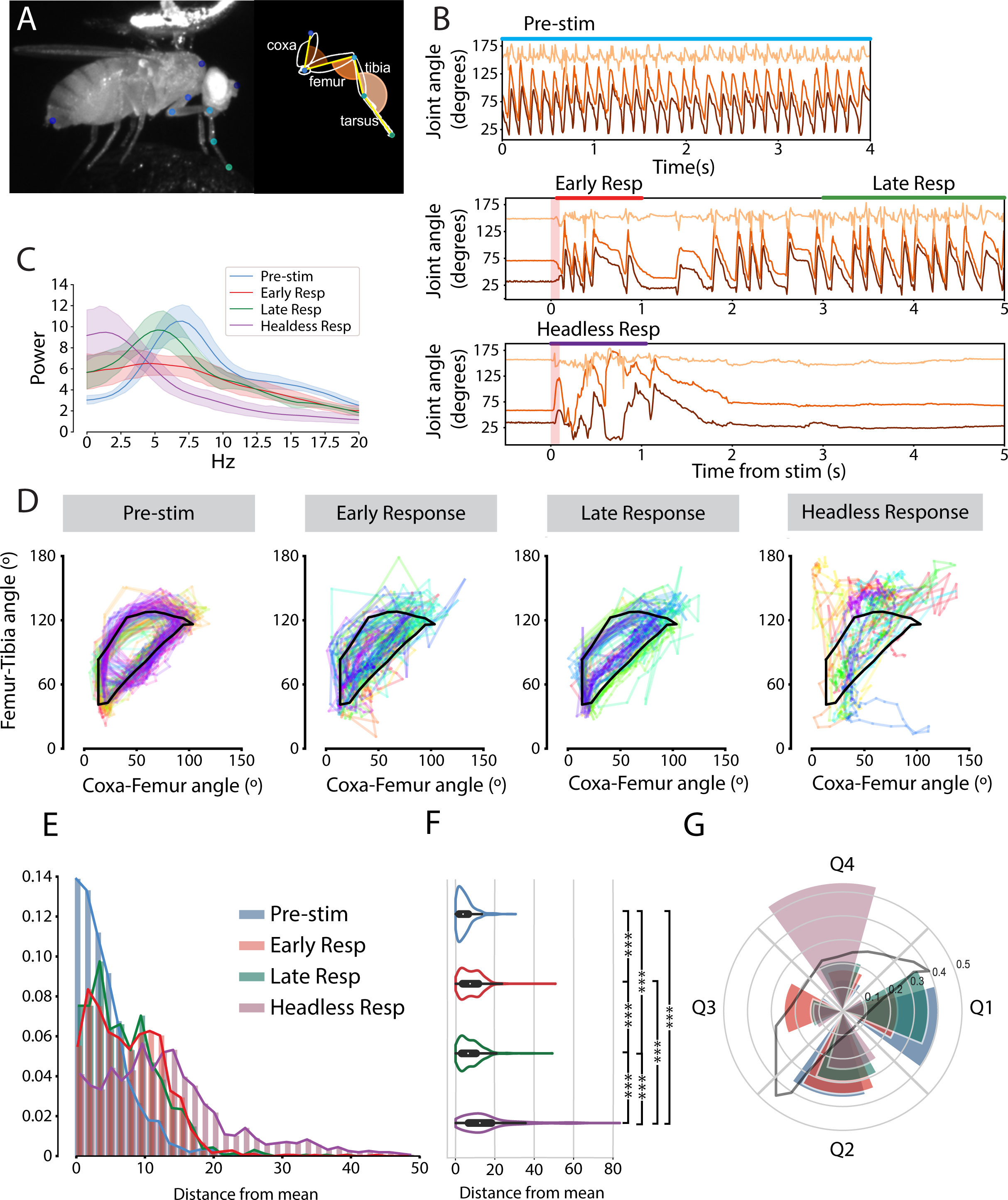
Mechanosensory Bristles evoked movement shows partially disrupted inter-joint coordination. (A) Tethered fly set-up showing tracked joint positions (blue/green circles). *Inset* schematic of Fly T1 leg showing 5 segments and 3 measured joint angles (Brown - Coxa-Femur angle, Bright Orange – Femur-Tibia angle, Light Orange – Tibia-Tarsus angle). (B) Modulation of joint angles throughout time for our 4 experimental conditions: Walking flies (blue horizontal bar), Head-Intact Early Response (red horizontal bar), Head-Intact Late Response (green horizontal bar) and Headless Early Response (purple horizontal bar). Red transparent vertical bars in the mid and lower panel indicate the time of optogenetic stimulation. (C) Fast Fourier Transform of leg angles for the Coxa-Femur Joint. (D) Angle-angle plots of Coxa-Femur vs Femur-Tibia angles over time. Different colors represent individual trials. Black line indicates the mean trace of the ‘walking’ condition over the step cycle. (E) Histogram of distances of points to the mean walking trace. (F) Violin plots for distances to mean walking trace. P values calculated using Dunns post-hoc test. ***P < 0.001 (G) Frequency of occupancy for quadrants of the Coxa-Femur angle vs Femur-Tibia angle space.

As expected, *pre-stimulation* walking flies showed stereotyped periodic patterns of joint angles (60) (Fig 6B). The Coxa-Femur joint and the Femur-Tibia joint were strongly modulated throughout the step-cycle in a correlated fashion, whilst the Tibia-Tarsus joint did not show an obvious correlation to the step cycle. We performed a Fast Fourier Transform (FFT) to extract component frequencies from the data (Fig. 6C). In *pre-stimulation* walking flies, the Coxa-Femur and Femur-Tibia joints showed the highest power at 7.0 Hz, corresponding to the average step frequency (Fig. 6C and Fig. S4A), while the Tibia-Tarsus joint was not periodically modulated (Fig. S4A). FFT analysis of the *early-response*, *late-response* and *headless response* also showed that Coxa-Femur and Femur-Tibia joints were more strongly modulated than the Tibia-Tarsal joint (Fig. S4A). We thus restricted our subsequent analysis to only Coxa-Femur and Femur-Tibia joints. To understand the regularity of motor behavior following stimulation we compared the FFT of our 4 conditions for the Coxa-Femur joint (Fig. 6C). Whilst in the intact-early stage the strong periodic modulation of joint angles seen in walking flies is lost, by the late stage, a peak has re-emerged (albeit around 5.2 Hz, a lower frequency than in the walking condition), showing the angles traces are once more becoming structured. Strikingly in headless flies, frequencies were highly variable with no clear peaks (Fig. 6C).

To further investigate the relationship between joints, we generated angle-angle plots between the Coxa-Femur joint and the Femur-Tibia joint for our 4 conditions (Fig. 6D). Angle-angle plots for walking flies showed a half moon-shaped ‘ring’ trajectory (Fig. 6D), corresponding to the stance and swing phases of the step cycle (Fig. S4B). Angle correlations were altered in the *early response* following stimulation (Fig. 6D). Whilst the half-moon outline was still visible, indicating that the fundamental relationship between joints has not been totally lost, the variability of the movement was greatly increased. Under *late response* conditions (i.e. 3-5 seconds after the stimulation) traces once more display a regular ‘walking-like’ ring (Fig. 6D). However, during *headless response* conditions, very distorted angle-angle traces are visible with an undetectable ring pattern. To quantify dispersion from the average walking cycle, we calculated the mean trace of the walking angles (Fig. 6D, black lines and Fig. S4B). We then analyzed the minimum distance for each measured point to the mean trace (Fig. 6E, F). Distances were significantly increased for all stimulated conditions as compared to pre-stimulus condition (Fig. 6F), indicating a significant disruption of the co-ordination between joints after MsB stimulation (P_walking-early response_ < 0.001, P_walking-late response_ < 0.001, P_walking-decapitated_ < 0.001, Dunn’s test). Distances were reduced from the *early response* to the *late response*, indicating a partial recovery of coordination (P_early response-late response_ < 0.001, Dunn’s test). In contrast, distances were greatly increased in the headless condition, indicating that loss of brain control leads to wildly disrupted relationships between joint angles (P_early response-decapitated_ < 0.001, P_late response-decapitated_ < 0.001, Dunn’s test). To understand which aspect of the relationship between angles was disrupted, we divided the angle-angle space into 4 quadrants (described in Fig. S4C). The positions of the lines were chosen to split the walking traces into 4 components of the step cycle – stance onset/quadrant 1 (corresponding generally to high Coxa-Femur angles); stance offset/quadrant 2 (corresponding to low Femur-Tibia angles); swing onset /quadrant 3 (corresponding to low Coxa-Femur angles); and swing offset/quadrant 4 (corresponding to high Femur-Tibia angles). We calculated the proportions of data points which fell in each quadrant for each condition (Fig. 6G). Pre-stimulus walking flies spend more time in quadrants 1 and 2, corresponding to the slow stance phase, and less time in quadrants 3 and 4 which correspond to the faster swing phase. Flies immediately after stimulation show an increase in occupancy of quadrant 3, likely corresponding to aborted swing initiations. Strikingly, headless flies show a large over-representation of points in quadrant 4, corresponding to extended Femur-Tibia angles, indicating that without descending control, extension is the default response, indeed to an extent where the flies almost invariably let the ball fall (Video S4). This effect was also observed in the gait-analysis experiments (Fig. 5 and Video S3).

Overall, our data show that inter-joint coordination is partially lost immediately after stimulation in head-intact flies, but has begun to recover by the period of 3 – 5 seconds after stimulation. In contrast, joint angles are highly uncoordinated following stimulation in headless flies, which show large overextensions of the Femur-Tibia joint, suggesting a role for descending control to appropriately regulate the relationships of joint angles to one another.

### Intra joint muscle coordination is maintained during movement triggered by MsB

Key for coherent locomotor activity is the reciprocal inhibition of opposing flexor and extensor muscles, each one controlled by their respective excitatory motor neuron pool (61–63). This implies that contraction of one joint muscle is concurrently matched by relaxation of its antagonistic muscle within the same joint. We asked if sensory-triggered movement also follows this muscle pattern, which we refer as intra-joint coordination. For this we monitored muscle contraction at the femur-tibia joint by measuring GCaMP fluorescence in leg muscles while optogenetically stimulating Leg MsB (Fig. 7A and Video S5) (64, 65), see Methods). The fly’s body and legs were glued to a coverslip (66), and muscle activity in the femur was recorded using a spinning disc confocal while the fly was stimulated with a 100ms light stimulus (see Methods). We identified the femur extensor and flexor muscles (Fig. 7B) and quantified the percentage of fluorescence in each frame, normalized to the maximum and minimum of the fluorescence for each video (Fig. 7B-D) (see Methods). In addition, we generated a kymograph covering both extensor and flexor muscles (Fig. S5). As expected in head intact flies, both controls groups (flies without stimulation and stimulated without all-trans-retinol (ATR)-supplemented food) display alternating muscle activity, meaning that antagonistic muscles were not active simultaneously (Fig. 7C, left and middle panel and kymograph on Fig. S5). We calculated the Spearman correlation between the two muscle signals over time, which generated negative values confirming their anti-phasic relation (Fig. 7E). Optogenetic stimulation of leg MsB showed a fast muscle response in accordance with the locomotor behavior previously observed. Interestingly, the observed muscle activity maintained the antagonistic alternation seen in control flies (Fig. 7C right panel, E and Fig. S5, Video S5). Stimulation of headless flies also led to an immediate muscle contraction showing an alternating antagonistic muscle activity, albeit with a higher variability (Fig. 7D, E, Video S5). Notably, both in head-intact and headless animals, we found no flexor or extensor muscle bias upon sensory stimulation (Data not shown), suggesting that no particular muscle was more responsive to sensory stimulation and that indeed a reciprocal motor pool inhibition mechanism is present to prevent muscle co-contraction upon stimulation (63, 67). We also compared the GCaMP fluorescence profile during muscle contraction by measuring peaks intensities and widths (Fig. 7F-H, see Methods). Noticeably, stimulation of leg MsB in headless flies led to a weaker muscle contraction pattern compared to head intact conditions (Fig. 7G), suggesting that higher order centers may act as modulators of motor neuron recruitment and activity. However, no differences were observed in the width of the muscle contraction peaks between head intact and headless flies (Fig. 7H).

**Figure 7.**
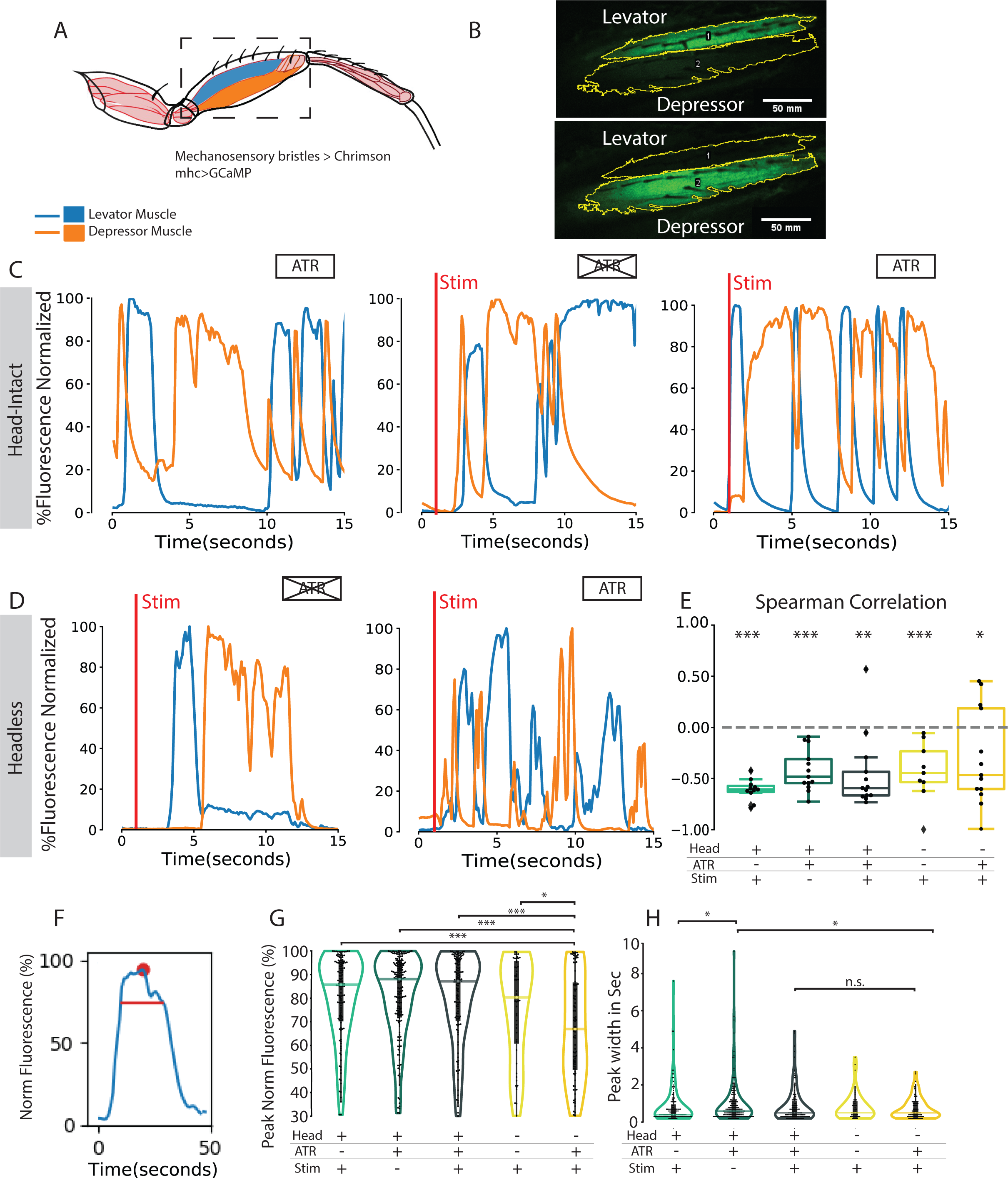
Muscle activity alternation is maintained during sensory-triggered movement. (A) Leg schematic showing tibia levator (blue) and depressor (orange) muscles in the femur. Adapted from (94). Dashed rectangle represents scanned region of the genotype: lexO-GCaMP6f; mhc-lexA:LHV2, dacRE-flp;Uas-FRT-stop-FRT-CsChrimson-mVenus, R65D12-Gal4 (B) Representative images of a femur showing levator (top panel) or depressor muscle (lower panel) contraction, resulting in GCaMP fluorescence inside the respective muscles. Yellow lines depict the ROIs used for fluorescence calculations. (C-D) Representative plots showing the fluctuation of fluorescence for levator (blue) and depressor (orange) muscle across time. Red bar represents the 100 ms red light stimulation. Values are normalized to maximum signal measured in each ROI throughout the experiment, corresponding to 100%. (C) Head-intact animals. Left panel: Non-stimulated animals in the presence of All-Trans Retinal (ATR) (N=11); Middle Panel: Stimulated animals in the absence of ATR (N=13); Right Panel: Stimulated animals in the presence of ATR (N=13) (D) Headless animals. Left panel: Stimulated animals in the absence of ATR (N=12); Right Panel: Stimulated animals in the presence of ATR (N=13). (E) Spearman correlation between levator and depressor muscle activity for each experimental condition. Positive values indicate that muscles are in-phase, while Negative values correspond to out of phase (antagonistic) activity. Boxplots represent the median as the middle line, with the lower and upper edges of the boxes representing the 25% and 75% quartiles, respectively, whiskers represent the range of the full data set, excluding outliers represented by diamonds. Filled dots represent individual flies. Statistical analysis with Kruskal–Wallis-ANOVA followed by Dunn’s *post hoc* analysis, no significant difference between the groups was detected. Wilcoxon test was employed to test if correlations were different from zero, all groups showed a significant negative correlation. (F) Schematic showing muscle contraction curve analysis shown in (G) and (H). Red dot indicates the maximum fluorescence value for the peak. Red line indicates the peak width at 20% below peak level. (G) Violin plot showing the maximum value of fluorescence for each contraction peak for each experimental condition. Each dot corresponds to a single muscle contraction peak. Horizontal line represents the median. (H) Violin plot showing the peak width for each contraction peak and each experimental condition. Each dot corresponds to the width of a single muscle contraction peak. Horizontal line represents the median. P values in (G, H) were calculated using Dunns post-hoc test. *P < 0.05; **P < 0.01; ***P < 0.001

Overall, our results show that leg MsB stimulation evoked movement largely maintains intra-joint muscle coordination properties in head-intact and headless flies.

## Discussion

In this study we took advantage of the sophisticated *Drosophila* genetic toolkit and kinematic analysis tools to understand mechanosensory evoked motor responses in different body regions. We found that a short 100-millisecond (ms) optogenetic stimulation of two different leg mechanosensory structures is sufficient to initiate forward movement in immobile animals (Fig. 1). In particular, we show that the motor response to MsB stimulation is dependent on the segment and number of cells being stimulated (Fig. 2 and 3). We found that MsB stimulation led a two-phase motor response comprised by an immediate *uncoordinated phase* followed by a late *coordinated phase* (Fig. 4; Video S3). While the coordinated phase shows all the fundamental features of normal locomotion, the uncoordinated phase lacks both inter and intra-leg coordination whilst sustaining antagonistic antiphasic muscle activity (Fig. 4, 6 and 7). Finally, the uncoordinated phase is mostly dependent on VNC circuits as leg MsB stimulated headless flies largely phenocopy the *uncoordinated phase* present in head-intact animals (Fig. 5, 6 and 7).

### Parallel activation of VNC Motor Circuits by sensory mechanosensory efferents

Studies using optogenetic approaches have shown the requirement of descending interneurons (DNs) in order to trigger forward and reverse locomotion (4, 5, 8, 68). In this study, we unveil a new parallel circuit for activation of VNC locomotor neural networks resulting in a fast, sustained forward motion albeit one defective in inter-joint and inter-leg coordination (Fig. S6). Indeed, we found a strikingly faster response upon MsB stimulation onset compared to ChO (peak at 60 ms after stimulus for MsB stimulation vs 600 ms after stimulus for ChO stimulation, Fig. 1D). It is noteworthy that previous studies have shown the generation of motor programs through the stimulation of sensory structures, independently of higher-order structures. For example, the mechanical and optogenetic activation of bristles in the entire body of the fly are able to elicit grooming movements (43), whereas the activation of specific bristles on the wing margin, which could be caused by an invading mite, leads to the initiation of a directed kick towards the stimulated area (17). These data highlight that the circuits required for the processing and execution of ethologically relevant behaviors are present in the VNC, which can be executed with remarkable speed and precision. Likewise, our data present evidence that neural networks in the VNC are capable of processing leg mechanosensory feedback to execute the appropriate cyclic motor program. Moreover, the location of the sensory input plays a major role in the selection of the motor outcome, instead of solely the class of the mechanosensory structures conveying the stimulus. Notably, stimulation of wing MsB did not induce uncoordinated walking but mostly grooming and kicking as previously described (17, 43) (Fig. 2). Since the leg and wing MsB target distinct neuropil regions in the VNC, namely the leg and wing neuropils (16) (Fig. S1), respectively, it is thus likely that distinct and possibly non-overlapping motor circuits become active.

Based on these data, we propose a model in which stimulation of MsB specifically in the legs activates VNC circuits in the leg neuropil that lead to a fast but uncoordinated motor output (Fig. S6, blue circuit). In parallel, the activity of MsB or other proprioceptive structures, perceive leg movements and relay this information, via ascending neurons to higher-order centers that inhibit uncoordinated movements and trigger coordinated motor output via descending interneurons (Fig. S6, orange circuit). The combined action of command DNs and MsBs can elicit a faster motor response, potentially allowing animals to efficiently escape physical danger or to avoid conspecifics under crowed situations (19).

### Sustained activity and segmental coupling

In this study, the movement evoked by the stimulation of MsB shows characteristic CPG-like activation, where antagonistic muscles maintain anti-phasic contraction leading to joint movement (69–72) (Fig. 7). The same behavior was observed in headless flies, implying a lack of interference of ascending or descending neurons in the sensory-evoked movement. These results are similar to the pharmacological stimulation of decapitated flies with stimulatory amines, notably serotonin, dopamine, and octopamine, which lead to locomotor bouts (73). Moreover, studies in deafferented VNCs using the muscarinic cholinergic agonist pilocarpine, have shown alternating and rhythmic activity in antagonistic leg motor neuron pools driven by the CPGs (69-72, 74-76). Despite this large body of work, the identity, location and architecture of the CPG population remains largely elusive. The reason for this gap in knowledge is the difficulty in accessing the neuronal VNC population, adding to the problems in separating the outputs from CPG and the activity in neurons that process sensory information during stepping. Nevertheless, some interneurons were identified as being part of this network in the cockroach and in the stick insect (27, 77). The use of mechanosensory stimulation in the fly provides an attractive port of entry in order to further identify and perform physiological characterization of leg CPGs taking advantage of the expanding *Drosophila* genetic toolkit (78). The second-order neurons from the mechanosensory fields remain elusive although electrophysiological data suggest tactile bristles establish synapses with intersegmental cholinergic and glutamatergic neurons, including ascending interneurons (13). Moreover, data from the locust suggest that these afferents target spiking local interneurons, some of which synapse on motor neurons (47). The reconstruction of serial-section electron microscopy (EM) datasets from the *Drosophila* VNC will provide a detailed connectome profile of the leg MsB (79).

### Cyclic activity and intersegmental coordination

Remarkably, we found reduced intersegmental coordination between hemi segments and leg joints immediately after MsB stimulation (Figs 4-6), albeit with antagonistic muscle contraction still being maintained (Fig. 7). These data resemble the effects of the muscarinic agonist pilocarpine in deafferented insect VNC preparations, which generate rhythmic activity of alternating motor neuron pools controlling joint movement (70, 75, 76, 80–82). Moreover, this rhythmic activity is independent within the three leg joints, as these rarely exhibit coordinated activity between them, meaning that different CPGs control the antagonistic activity of motor neuron pools of each leg joint (72, 74, 83, 84). Given that in our experimental design, animals still maintain their proprioceptive features that assist in segment coupling, we can speculate that the MsB-dependent cyclic activity overrides or is insensitive to sensory feedback or to any other coupling mechanism. Our data also suggest a role for descending feedback promoting segmental coordination since decapitation decreased the degree of inter-and intra-leg coordination (Figs. 3F, G and 4F, S6). Work on hemimetabolous insects has shown a role for descending neurons (DNs) that traverse the neck controlling the onset and features of walking patterns (reviewed by (74, 85)). Moreover, optogenetic manipulation of defined populations of DNs can trigger stereotyped behaviors and change walking patterns (5, 86). Multisegmental coupling is provided either by sensory feedback, descending inputs or centrally by nonspiking interneurons (72, 87). The relevance of each component depends largely on the speed the animal typically assumes, with slow-walking animals - such as the stick insect - relying heavily on sensory feedback (26, 88, 89). Regardless, our results suggest that, as in other insects, alternating activity of antagonistic motor pools in *Drosophila* are uncoupled between different joints and hemisegments, further reinforcing the use this model system to unravel the neuronal architecture responsible for coordinated motor patterns.

## Methods

### Fly strains

All flies used in this study were non-virgin females between 1 and 7 days post-eclosion. Flies were housed in a 25° incubator with 50-70% humidity. Prior to experimentation, all flies were manipulated under cold anesthesia and allowed to recover for at least 24h. The GAL4 lines from the Janelia Flylight Collection were: R65D12-Gal4(attP2), R77E09-Gal4(attP2), R85A07-Gal4(attP2), R45D07-Gal4(attP2), R54A11-Gal4(attP2), R39A11-Gal4(attP2) and R64D07-Gal4(attP2), R50A05-Gal4(attP2), R22E04-Gal4(attP2), R79E02-gal4(attP2) and pBDP-GAL4UW (w[1118];P{GAL4.1Uw}attp2) (EmptyGal4). Ato1.9-flp and Vg^BE^-flp were generated by TOPO cloning into an entry clone the 1.9kb enhancer fragment from the regulatory region of atonal (90), and the boundary enhancer of vestigial (Vg^BE^) (91, 92) (a gift from Mirian Zecca), respectively. Both entry clones were subjected to a Gateway reaction into a flp destination plasmid. Transgenic lines were generated by standard procedures in a *yw* background. The dac^RE^-flp was used in previously (11). UAS-FRT-stop-FRT-CsChrimson-mVenus was a gift from Gerry Rubin (39). UAS-FRT-stop-FRT-GFP, Lexop-GCaMP6f and ey-flp lines were obtained from the Bloomington Stock Center. For the calcium imaging in leg muscles we used lexO-GCamp6f/;mhc-lexA,lexO-Cherry/SM6^Tm6b (65) and the recombinant line Dac^RE^-flp;Uas-FRT-stop-FRT-CsChrimson-mVenus (VK5), R65D12-Gal4 (attP2).

### Behavior Setup and optogenetic activation of mechanosensory structures

The open field behavior arena for optogenetic stimulation was custom built from 3 mm opaque white and black acrylic sheets using a laser cutter. The circular arena had a diameter of 54mm, and was surrounded by 6 deep 655 nm red Leds (LUXEON Rebel Led’s) evenly distributed providing a stimulation power of 0.016 mW/mm^2^ (Fig. 1C), The edge of the arena was surrounded by water to prevent flies accessing the walls. The top of the arena (transparent acrylic) was covered with Sigmacote (Sigma-Aldrich). A white LED array was placed under the arena to serve as backlight. Flies were imaged from above using a camera (Flea3-USB3, Point Grey, Teledyne FLIR LLC) equipped with a 12 mm Computar (Commack, NY) lens covered with a 600nm, OD 4.0 Shortpass Filter (Edmund optics, Barrington, NJ) to block optogenetic stimulation light.

For the experiments in Figure 1, we used 1-7 day old females, comprising 20 control flies (empty-GAL4), and 20 flies expressing CsChrimson in leg mechanosensory bristles. Flies were bred and kept in food infused with all trans-Retinal (Final Concentration 0.4 mM; R2500, Sigma-Aldrich). Single flies were aspirated into the chamber. We waited for a spontaneous pause in behavior (no walking or grooming) and manually delivered a single 1-pulse stimulation of 100ms. Imaging continued for at least 20 seconds after stimulation. For decapitated fly experiments, flies were anesthetized on a cold plate and their head was cut off. After the surgery flies were kept in an empty vial for 1-5 minutes prior to testing.

### Video acquisition and Tracking

The video image was acquired at 30 Hz (1200 x 1210) using Fly capture software. Image segmentation was performed by custom software in python using OpenCV. Two main features were extracted from the videos: fly position (X and Y coordinates) and pixel change around the fly. Positions were calculated from the centroid of an ellipse fitted to the fly by background subtraction. The pixel change was quantified by calculating the number of active pixels in a 100×100 pixel region surrounding the centroid. A pixel was considered to be active if it recorded a change higher than 10 intensity levels.

The fly distance and speed were calculated for each frame using fly position x and y coordinates with the formula:

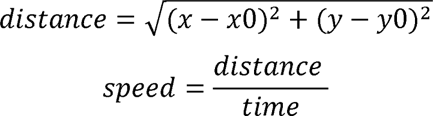

### Behavioral Classifiers

We used velocity and pixel change features to automatically classify behavioral states. We classified fly behavior into 4 categories: Walking, Grooming, Stopping and Leg Displacement. A fly is considered to be walking if its average speed is higher than 1.9mm/sec. Grooming is detected when the velocity of the fly is lower than 1.9 mm/s and average pixel change of a section of 8 frames is above 180 pixel/s. We classified as leg displacement the movement in which velocity is lower than 1.9 mm/s and average pixel change of a section of 8 frames is between 30 and 180 pixel/s. Finally, a fly is considered to be stopping when velocity is lower than 1.9 and pixel change is lower than 30 pixel/s. Thresholds were determined and confirmed by manual annotations of fly behavior (% of efficacy). Later, walking behavior was either classified into coordinated or uncoordinated walking by visual inspection (Fig. 2 and 3).

### fTIR imaging and FlyWalker software

For the analysis of walking behavior upon stimulation of sensory structures a new arena comprising 12 LEDs (Deep red 655nm, Luxeon Rebel LED) was incorporated in the previously described fTIR setup (11), rendering a stimulation power of 0.014 mW/mm^2^. Flies were placed in the FlyWalker arena and a 100 ms pulse of red light was given. Each fly only received 1 pulse. The walking kinematics of flies after stimulation were quantified using FlyWalker software package. One more parameter called Non-Compliance Index was added to the previously published Kinematics parameters. Non-Compliance Index is defined as the fraction of frames in a video in which the fly displays non-compliant leg combinations. Non-canonical stance combinations correspond to patterns that do not fit either of the idealized tripod, tetrapod and wave gaits (11, 36). In the experiments where we used headless flies, we glued the head back onto the thorax using UV curable glue (Loctite 3104, Henkel Adhesives) in order to avoid alterations in the body weight that could potentially change walking patterns of the fly and to facilitate tracking.

### Tethered fly imaging

For imaging of leg angles in Figure 4 we used 3-7 days-old females expressing CsChrimson in mechanosensory bristles, as above. Adult flies were kept in food infused with all trans-Retinal since the day of eclosion (Final Concentration 0.4 mM; R2500,Sigma-Aldrich). Flies were anaesthetised on ice, and either decapitated with scissors (Fine Scientific Tools) or left with head intact. They were positioned on a cold plate with the thorax upwards, and a wire was glued to the thorax with UV curable glue (Norland Optical Adhesives #81). Flies were fixed in the tethered set-up using the wire above a porexpam ball (4.5 mm diameter, painted with black paint (Posca Uni)), held in a custom made holder (the glass funnel of a Pasteur pipette). Flies could freely pick up and walk on the ball, and if dropped could regain it again. Each fly was allowed to recover for 5 minutes on the set-up. After 5 minutes we waited for a spontaneous pause in behavior and delivered a 100ms light stimulus via a single deep red LED (wavelength 627nm), positioned 3 cm away from the fly. Light power at the fly was 13.8 nm/cm^2^, calibrated to produce a motor response in CsChrimson flies whilst not affecting flies in food lacking all trans-Retinal (data not shown). Animals were illuminated with an InfraRed LED (Thorlabs) and imaged with a PointGrey Camera (Flea3) with a Computar Macro-zoom lens and a long-pass filter, at a frame rate of 120 fps. Video capture and LED stimulation were controlled using a custom built Bonsai workflow (93).

Walking bouts were extracted from head-intact non-stimulated flies (either before stimulation or in sessions where no stimulation was delivered), and were manually classified as satisfying straight walking with at least 6 step cycles. In total 8 walking bouts were used from 3 flies. In the Head-intact and headless stimulation conditions for the ‘early response’ and ‘late response’ timepoint, we visually inspected videos and excluded timepoints where the flies were either still or performing grooming, as being non-informative for our question of limb coordination while walking. Total N was 11 (head intact early response), 6 (head intact late response) and 10 (decapitated early response). Videos were analyzed using DeepLabCut (59). 18 points were tracked in total; 3 body points (head, thorax and tail) and 5 points per leg (Body-Coxa; Coxa-Femur; Femur-Tibia; Tibia-Tarsus; and Tarsus-end); however only T1 limb tracking was used for the final analysis. Limb angles for T1 legs were calculated from tracked positions using python.

### Leg and VNC dissection and Mounting

To observe the morphology of leg sensory neurons, legs were dissected from the body in PBS. The legs were then fixed in PFA 4% overnight at 4°C. After fixation legs were washed 3x in 0.3% Triton-X in PBS for 20 minutes at R.T and mounted in 70% glycerol. Brains and VNCs were fixed in PFA 4% for 20 minutes and dissected in PBS. Tissues were labeled with a primary antibody raised against brp (mouse nc82, 1:10; DSHB) followed by a secondary anti-mouse 647 (1:500; Jackson immunologicals). Tissues were mounted in in 70% glycerol and scanned in a confocal microscope Zeiss (Oberkochen, Germany) LSM710 confocal microscope with a 40× objective.

### *In vivo* calcium imaging during leg stimulation

Flies containing the following genotype: lexO-GCamp6f;mhc-lexA,lexO-Cherry and dacRE-flp;Uas-FRT-stop-FRT-CsChrimson-mVenus(VK5), R65D12-Gal4(attP2) were anesthetized with ice for 3-5 min. Three ipsilateral legs were glued to a glass slide with UV-Curable glue. The body of the fly was immobilized using beeswax. GCaMP imaging was performed at 25°C on a Nikon/Andor Revolution XD spinning-disk confocal microscope with an EMCCD camera (iXon 897) using iQ 3.5 software and using a 20× CFI Super Fluor 20X 0.75 NA dry objective. Images were taken at the scan speed of 100ms per frame. A set of 6 deep red LEDs was paced around the fly to provide the 100ms red light pulse.

### Data processing and Analysis

We extracted motor parameters using FlyWalker Software. Since many of the measured gait parameters vary with speed, we analyzed the data for these parameters by determining the best-fit regression model of each individual parameter with speed for the control experiment and then determining the residual values for each experimental group in relation to this regression model. Data was then expressed as the difference to the residual normalized line. Normalized data was previously tested for normality and homoscedasticity with Shapiro-Wilk and Levene tests. Statistical differences between experimental groups were determined using Kruskal-Wallis analysis of variance followed by Dunn’s post hoc test (for non-normal distributions) or one-way-Anova followed by Tukey’s post hoc test (for normal distributions). Data analysis was performed using custom Python scripts. Boxplots represent the median as the middle line, with the lower and upper edges of the boxes representing the 25% and 75% quartiles, respectively. Whiskers represent the range of the full data set, excluding outliers. Outliers are defined as any value that is 1.5 times the interquartile range below or above the 25% and 75% quartiles, respectively. For the boxplots in fig 4 and 5 we decided to use raw data not normalized to the control since Non-Compliance index and Stance straightness parameters do not show a correlation with velocity (data not shown). To have a more succinct representation of the data we used Principal Component Analysis (PCA), an unsupervised dimensionality reduction method. PCA finds a linear projection of the data from a high-dimensional space onto a lower-dimensional subspace, while maximizing variance of the projected data, and thus retains meaningful information, resulting in a description of the data as a function of a smaller set of uncorrelated variables (or principal components). The data were first pre-processed by centering and scaling, after which the PCA algorithm computed the covariance matrix in order to determine the correlation between variables and calculated the eigenvectors and eigenvalues of the covariance matrix in order to identify the principal components. We chose the first three principal components to visualize the data. The first two components were chosen to generate a two-dimensional plot. Each dot in the plots corresponds to a video, representing these new abstract variables. Color-coded dots were used to distinguish specific groups. As such, clusters of dots reflect similar walking patterns, shared by the corresponding flies (48).

For the analysis of the muscle fluorescence, we used Fiji software. To delineate the ROI Area of the depressor and levator muscle we used the threshold tool and measured the mean fluorescence value for each frame using ROI Manager, multimeasure tool. Fluorescence values for each muscle were normalized by subtracting the minimum value of each video and dividing by the maximum value, giving contraction values between 0 and 100%. Muscle contraction peaks were detected with a peak finding algorithm in Python. Peak width was calculated by subtracting fluorescence 20 percentage points from each peak value, and measuring the width of the signal that exceeded this value around each peak (Fig 7F).

## Supporting information

Supplementary Figure 1. Brain and VNC Expression patterns

Supplementary Figure 2. Cumulative distance in flies exposed to sensory stimulation

Supplementary Figure 3. Validation of behavioral classifier

Supplementary Figure 4. Fast Fourier Transform and Quadrant analysis of leg angles

Supplementary Figure 5. Kymographs of Muscle GCaMP fluorescence

Supplementary Figure 6. Proposed model for MsB - evoked motor response

## Acknowledgements

We thank Mendes lab, Rita Teodoro and her lab for comments and suggestions during the execution of this project, Allan Mancoo for feedback on data analysis, Inês Fernandes for experimental assistance, Rita Fernandes for advice on Figure 6, and Luísa Vasconcelos, Turgay Akay and Daniela Pereira for comments on the manuscript. We thank Mirian Zecca and Richard Mann for support generating the transgenic lines used in this study. We also thank CONGENTO: consortium for genetically tractable organisms, the Fly Platform and the Scientific Hardware Platform at the Champalimaud Centre for the Unknown, and the Bloomington Drosophila Stock Center for fly stocks. This work was supported by Fundação para a Ciência e a Tecnologia (FCT) (PTDC/BIA-COM/0151/2020), iNOVA4Health (UIDB/04462/2020 and UIDP/04462/2020), and LS4FUTURE (LA/P/0087/2020) to CSM. AM was supported by a doctoral fellowship from FCT (PD/BD/128445/2017).

## Author contributions

AMM, AFH and CM contributed to conception and design of the study. AMM performed the experiments and analyzed the raw data; AFH performed data acquisition and analysis of Figure 6, and contributed to the analysis of Figure 7 data. GB contributed for analysis and data acquisition of Figure 3. AMM, AFH, MM and CSM wrote the manuscript. All authors contributed to manuscript revision, read, and approved the submitted version.

## Supplementary Information

**Supplementary Figure 1. Brain and VNC Expression patterns**

(A-D) Green: mVenus fluorescence; Magenta: anti-Bruchpilot. Top panel: central brain; Lower panel: VNC. Bar, 100 μm. In (A-B) middle Panel: Anterior view of Protothoracic Neuromere (ProNm) (A) Leg MsB1 (Genotype: R65D12-GAL4,DAC^RE^-flp,UAS>> csChrimson-mVenus).(B) Leg femoral ChO (Genotype: R79E02-GAL4, DAC^RE^-flp, UAS>> csChrimson-mVenus). (C) Wing MsB (Genotype: R79E02-GAL4, vg^BE^-flp, UAS>>csChrimson-mVenus). (D) Eye MsB (Genotype: R79E02-GAL4, ey-flp, UAS>> csChrimson-mVenus). (E) Wing expression pattern for wing MsB line (Genotype: R79E02-GAL4, vg^BE^-flp, UAS>>csChrimson-mVenus). Arrows indicate cell bodies of MsB. Magenta: cuticle auto-fluorescence. Bar, 100 μm.

**Supplementary Figure 2. Cumulative distance in flies exposed to sensory stimulation**

Cumulative distance over time before and after 100 ms red light stimulation (red bar) in (A) head-intact and (B) headless flies for the Empty Control group (Left panel), MsB1 lines (middle panel) and leg ChO line (right panel).

**Supplementary Figure 3. Validation of behavioral classifier**

(A)Scatter plot depicting speed and pixel change of bouts of immobility (beige), Walking (orange), leg movement (brown) and grooming (green). Each data point corresponds to 1 frame, n >= 250 for each behavior. Vertical and horizontal lines indicate the thresholds used to classify each behavior. Walking (>1.9mm/s), grooming(<1.9mm/s and >180 pixel/s), leg movement (<1.9mm/s and 30-180 pixel/s), immobility (<1.9mm/s and <30pixel/s) see methods. (B) Percentage of accuracy of the behavioral classifier for each behavior. (C) Duration of movement after a single pulse of red light stimulation (100ms) for each MsB line. Lines are ordered according to the leg segment in which they have more expression, from proximal to distal.

**Supplementary Figure 4. Fast Fourier Transform and Quadrant analysis of leg angles**

(A) Fast fourier transforms of traces for Coxa-Femur, Femur-Tibia and Tibia-Tarsus angles for the 4 different experimental conditions (Walking, Early Response, Late Response, Headless). (B) Walking leg angles divided into the swing and stance phases. Black line – mean trace. Arrowheads – direction of movement as the fly progresses through a step cycle. (C) Angle-angle plots divided into 4 quadrants. Quadrant lines were calibrated to divide the Walking trace into 4 component parts: Stance Early (Right Quadrant), Stance Late (Lower Quadrant), Swing Early (Left Quadrant), Swing Late (Upper Quadrant).

**Supplementary Figure 5. Kymographs of Muscle GCaMP fluorescence**

(A) Representative image of fly femur depicting the ROI used to measure fluorescence in levator and depressor muscles. (B) Kymographs of fluorescence corresponding to the ROI in (A) for head-intact animals: Non-stimulated in the presence of All-Trans Retinal (ATR), stimulated in the absence of ATR and stimulated in the presence of ATR; for healdess animals: stimulated in the absence of ATR and stimulated in the presence of ATR. Red Bar represents the frame where the 100 ms red light pulse was given.

**Supplementary Figure 6. Proposed model for MsB - evoked motor response**

Proposed model representing the possible mechanisms behind the motor output triggered by MsB stimulation. Stimulation of MsB in the legs activates VNC circuits in the leg neuropil that lead to an initial uncoordinated motor output (blue). Simultaneously, ascending neurons, possibly activated by either MsB or other proprioception structures, sense leg movement and relay this information to higher-order centers that inhibit uncoordinated movement and trigger coordinated motor output via descending interneurons (orange). Both uncoordinated and coordinated motor output enable a rapid locomotor response.

**Supplementary Table 1**

Cell labelling count for each leg segment for the 7 MsB lines.

**Supplementary Video 1**

Activation of MsB and ChO in Head intact and Headless flies in open arena.

**Supplementary Video 2**

Activation of MsB in the leg, wing or head of head-intact and headless flies in open arena.

**Supplementary Video 3**

Tracking of leg movement using FlyWalker system, after stimulation of MsB in Head-Intact and Headless Flies

**Supplementary Video 4**

Tracking of leg movement in tethered flies walking on a polystyrene ball, after MsB stimulation. Four conditions were assessed: walking flies (pre-stimulation), head-intact flies early response (1s after stimulation onset) and late response (3-5s after stimulation onset), headless flies (1s after stimulation).

**Supplementary Video 5**

GCaMP activity of tibia Levator and Depressor muscles in the femur of the fly. Three conditions were depicted: Stimulation of MsB in Head-Intact flies reared without ATR; Stimulation of MsB in Head-Intact flies reared with ATR; Stimulation of MsB in Headless flies reared with ATR.

