## Supplementary figures and images for "Mechanosensory stimulation triggers sustained local motor activity in *Drosophila melanogaster*"

### Supplementary Figure 1. Brain and VNC Expression patterns

#### Supplementary Fig 1

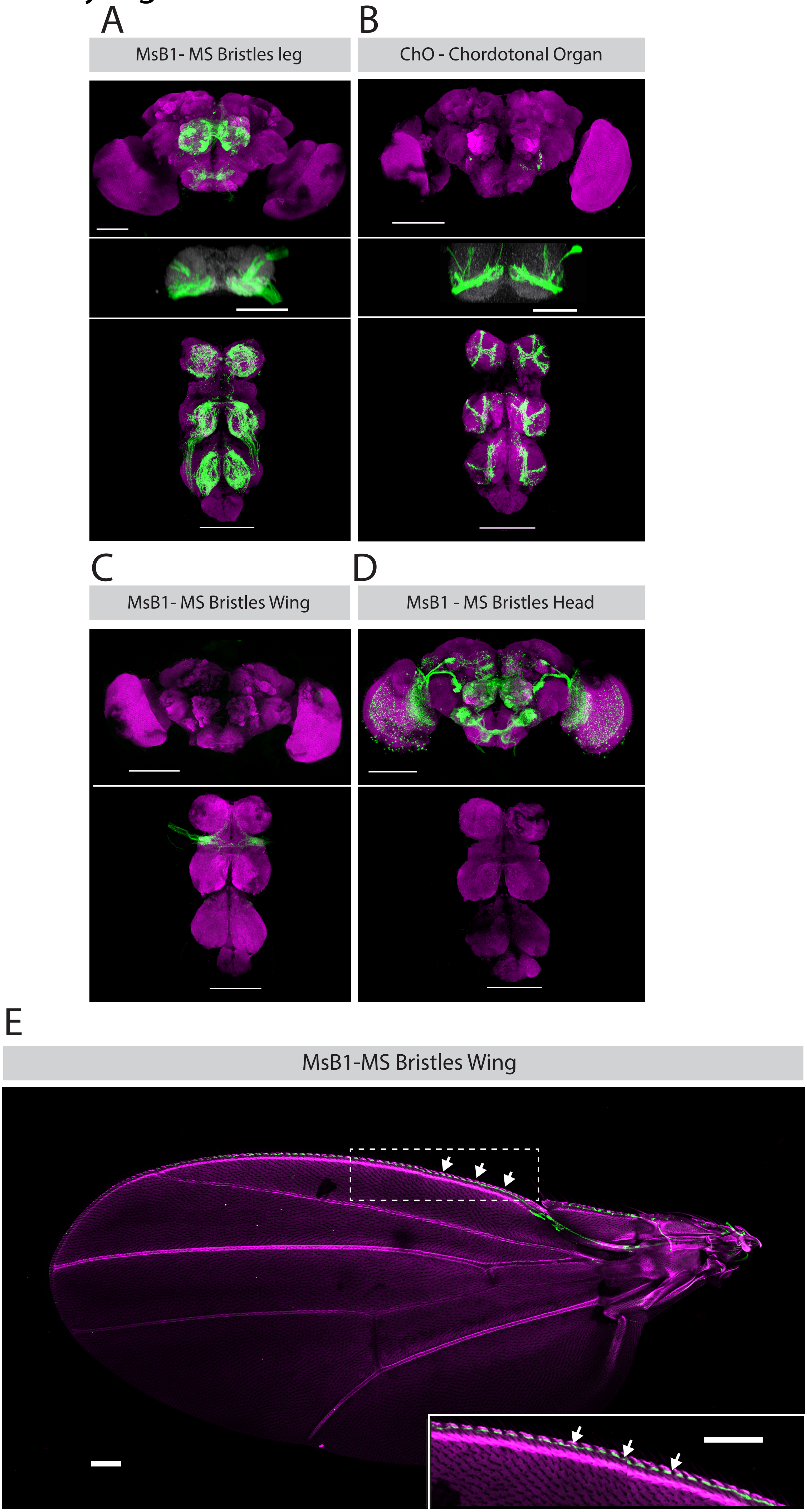

### Supplementary Figure 2. Cumulative distance in flies exposed to sensory stimulation

#### Supplementary Figure 2

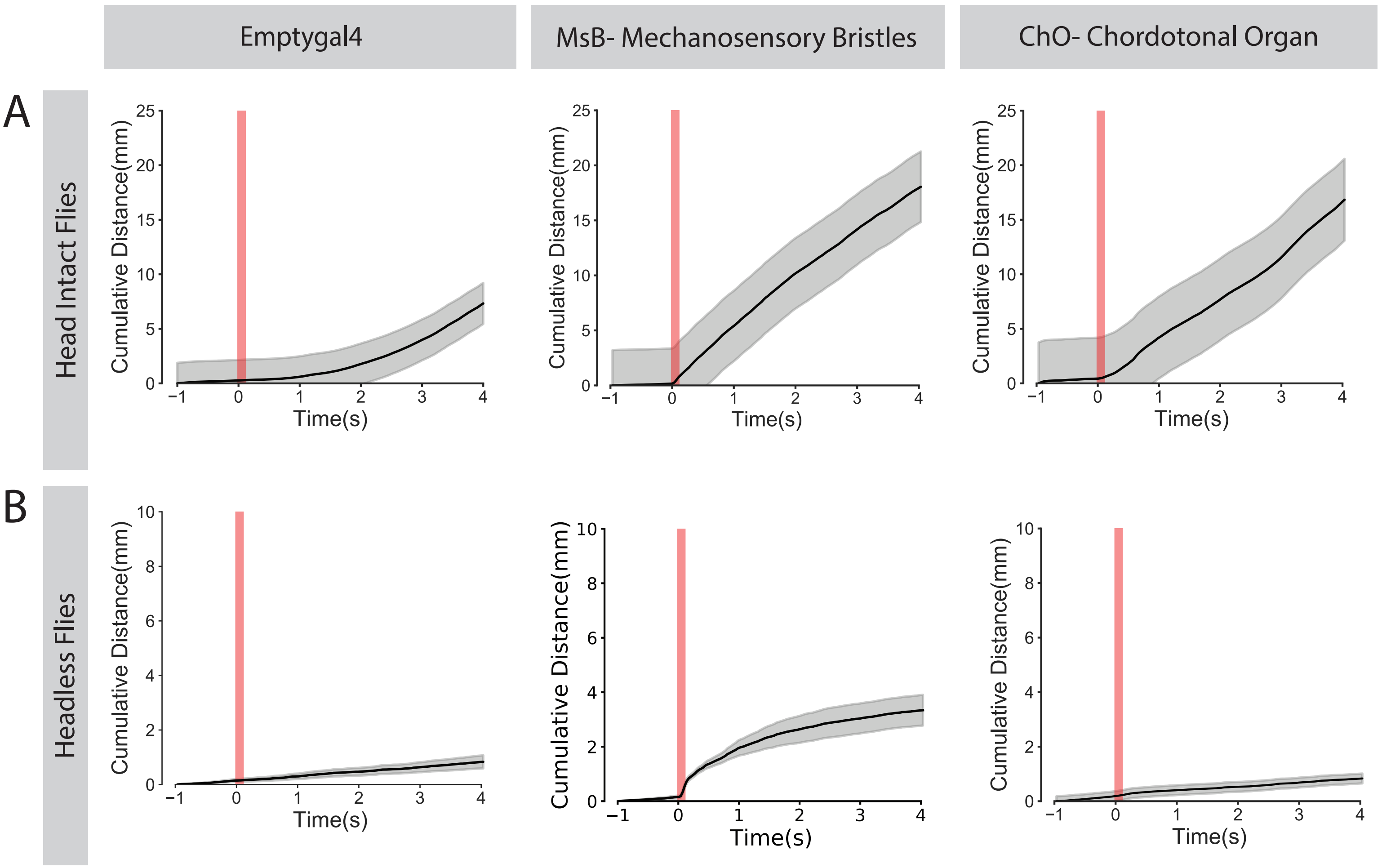

### Supplementary Figure 3. Validation of behavioral classifier

#### Supplementary Figure 3

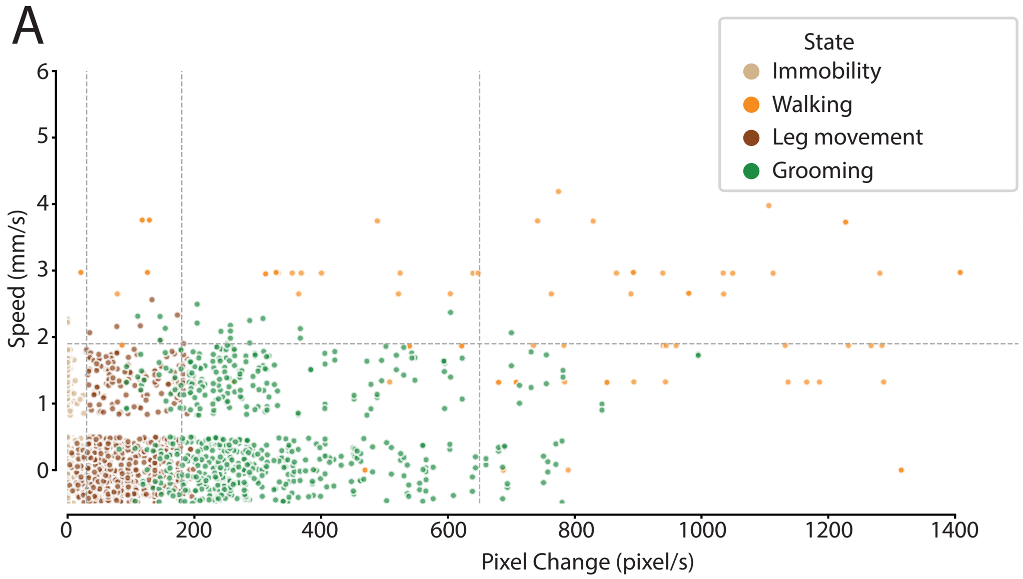


B

| Type of Movement | Accuracy |
|---|---|
| Immobility | 99.09% |
| Walking | 91.54% |
| Leg movement | 90.15% |
| Grooming | 82.87% |

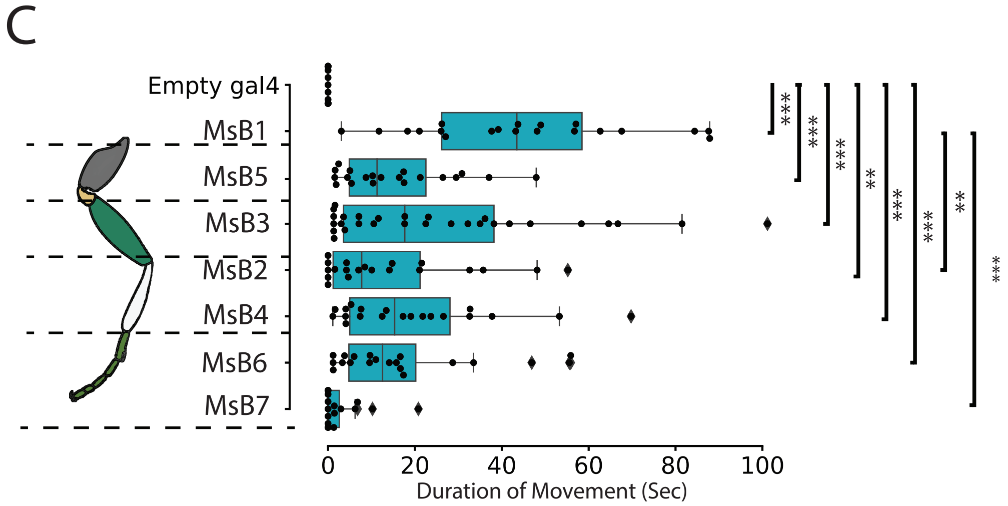

### Supplementary Figure 4. Fast Fourier Transform and Quadrant analysis of leg angles

#### Supplementary Fig 4

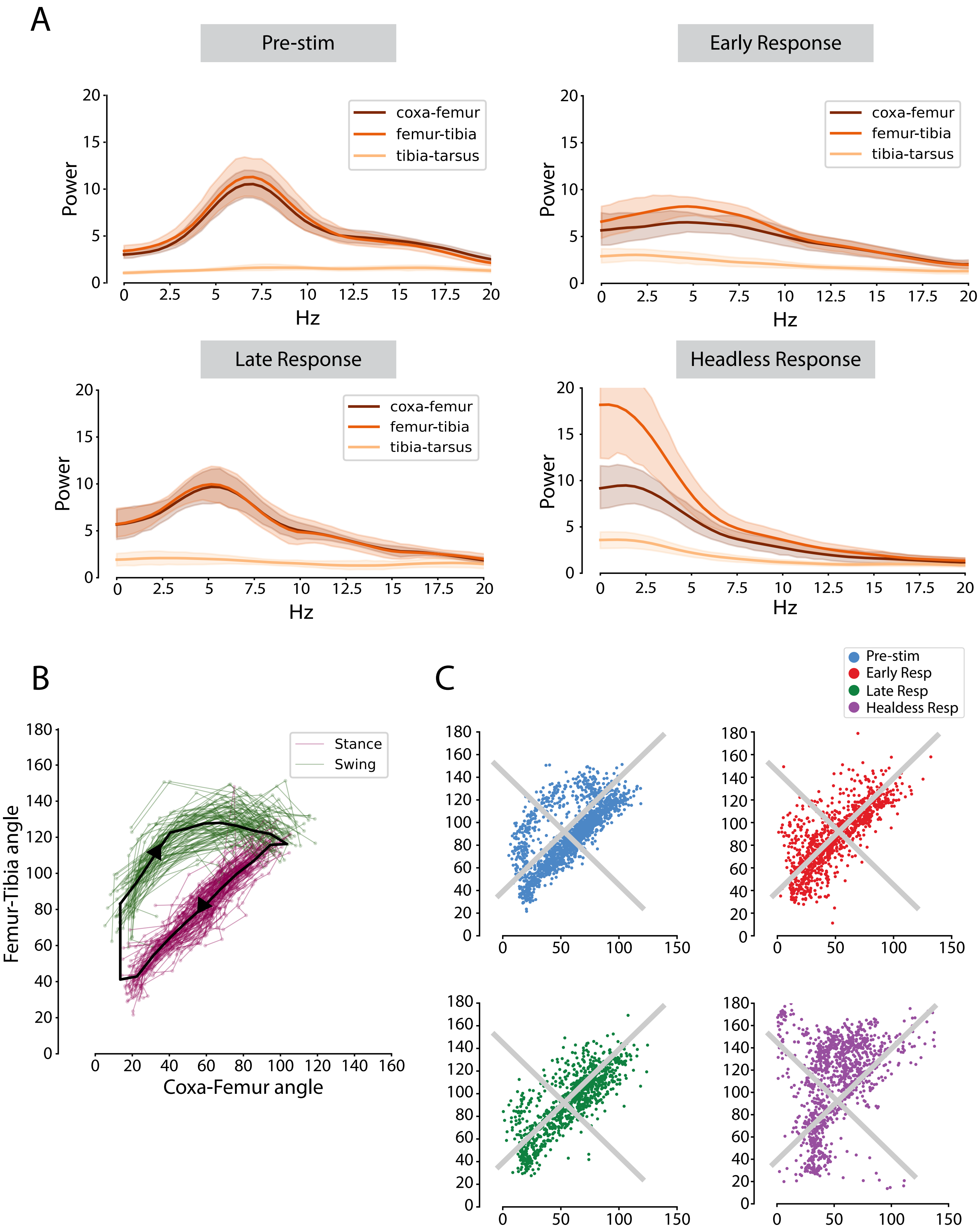

### Supplementary Figure 5. Kymographs of Muscle GCaMP fluorescence

#### Supplementary Fig 5

A

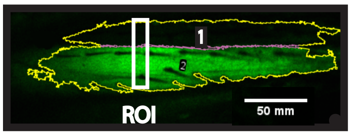


B

| Head | ATR | Stim |
|---|---|---|
| + | + | - |
| + | - | + |
| + | + | + |
| - | - | + |
| - | + | + |

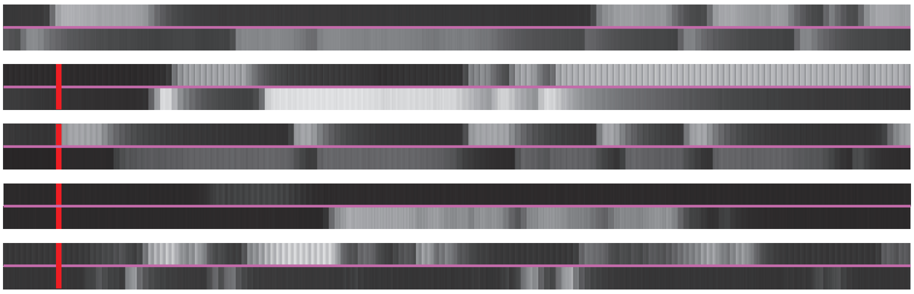

### Supplementary Figure 6. Proposed model for MsB - evoked motor response

#### Supplementary Figure 6

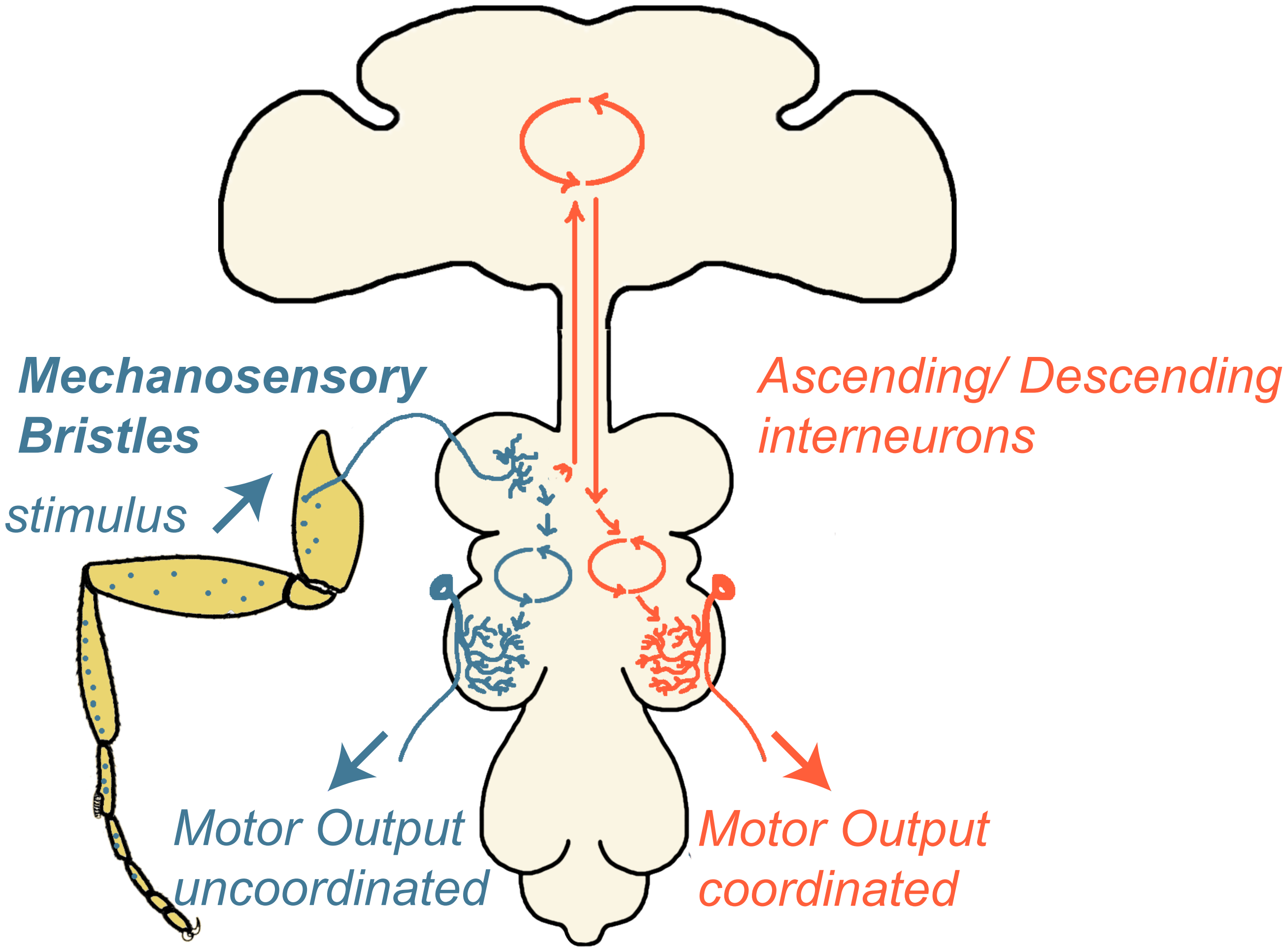
