## Supplementary Table 1 for "Mechanosensory stimulation triggers sustained local motor activity in *Drosophila melanogaster*"

Table 1. Cell labelling count for each leg segment for the 7 MSB lines.

| | Coxa | Trochanter | Femur | Tibia | Tarsus | Total | N |
|---|---|---|---|---|---|---|---|
| MSB1 | 5±1.26 | 7±0.63 | 37.75±4.43 | 48.25±11.61 | 70.75±10.18 | 168.75±15.2 | 4 |
| MSB2 | 0 | 0 | 0 | 50.5±7.23 | 6±4 | 56.5±9.6 | 4 |
| MSB3 | 0 | 0 | 41.42±5.42 | 4.86±1.45 | 1.29±.48 | 47.57±7.11 | 7 |
| MSB4 | 0 | 0 | 1±1.22 | 32±12.9 | 10.25±3.49 | 43.25±12.17 | 4 |
| MSB5 | 2.25±1.60 | 10.5±2.05 | 14.25±4.49 | 8.25±3.42 | 7.25±4.76 | 42.5±7.63 | 4 |
| MSB6 | 0.5±0.77 | 0 | 1.5±2.06 | 0.5±0.5 | 23.75±2.49 | 26.25±3.86 | 4 |
| MSB7 | 0 | 0 | 0 | 0 | 4.25±0.43 | 4.25±0.43 | 4 |
